# Molecular recording of calcium concentration via calcium-dependent protein proximity labeling

**DOI:** 10.1101/2022.07.14.500122

**Authors:** J. Wren Kim, Adeline J. H. Yong, Erin E. Aisenberg, Wei Wang, Ted M. Dawson, Valina L. Dawson, Ruixuan Gao, Helen S. Bateup, Yuh Nung Jan, Nicholas T. Ingolia

## Abstract

Calcium ions serve as key intracellular signals. Local, transient increases in calcium concentrations can activate calcium sensor proteins that in turn trigger downstream effectors. In neurons, calcium transients play a central role in regulating neurotransmitter release and synaptic plasticity. However, it is challenging to capture the molecular events associated with these localized and ephemeral calcium signals. Here we present an engineered biotin ligase that combines the power of genetically encoded calcium indicators with protein proximity labeling. The enzyme, Cal-ID, biotinylates nearby proteins within minutes in response to elevated local calcium levels. The biotinylated proteins can be identified via mass spectrometry and visualized using microscopy. In neurons, Cal-ID labeling is triggered by neuronal activity, leading to prominent protein biotinylation that enables transcription-independent activity mapping in the brain. Cal-ID produces a biochemical record of calcium signaling and neuronal activity with high spatial resolution and molecular specificity.

## Introduction

Calcium ions (Ca^2+^) are universal second messengers in eukaryotic cells. Ca^2+^ play essential roles in cellular physiology and Ca^2+^ homeostasis is critical to cell viability. Cytosolic Ca^2+^ levels are typically maintained at ∼100 nM, which is more than 10,000-fold lower than the extracellular Ca^2+^ levels^1^. However, cytosolic Ca^2+^ can increase rapidly due to release from intracellular stores or influx through plasma membrane channels. The resulting Ca^2+^ transients activate Ca^2+^ sensor proteins, which often bind to effector proteins in a Ca^2+^-dependent manner to exert regulatory function. Ca^2+^ signaling is often localized to subcellular microdomains that form near the sites of Ca^2+^ influx as well, and thus can control both the position and timing of downstream responses^2^.

Ca^2+^ signaling features prominently in neuronal physiology, where tight spatial and temporal control of signaling events is critical. Ca^2+^ regulates neurotransmitter release at the presynaptic active zone, and neurotransmitter receptors in turn allow Ca^2+^ influx into the postsynaptic region; additionally, G-protein-coupled neurotransmitter receptors trigger endoplasmic reticulum (ER)-mediated Ca^2+^ signaling^3^. Local Ca^2+^ dynamics play a key role in synaptic plasticity, and activity-dependent gene expression requires Ca^2+^ signaling to the nucleus^4^.

Indeed, Ca^2+^ dynamics can serve as a valuable proxy of neuronal activity. Voltage-gated Ca^2+^ channels allow Ca^2+^ influx from the extracellular space when the plasma membrane depolarizes^5^. Therefore, chemical and genetic tools to monitor Ca^2+^ levels, such as Fura and Fluo dyes, and genetically encoded Ca^2+^ indicators such as GCaMPs, have been extensively utilized to report on neuronal activity in real time^6^. These tools, which enabled researchers to track activities of many neurons via live imaging, have become a landmark technological advance in neuroscience. However, they do not provide information about the molecular events occurring at the site of Ca^2+^ signaling events.

In the last decade, proximity labeling has emerged as a new approach for surveying macromolecular interaction and localization in living cells^7^. In proximity labeling, an engineered enzyme covalently modifies nearby proteins, often by generating diffusible but short-lived reactive compounds. For example, the *E. coli* Bifunctional ligase/repressor (BirA) has been adapted to promiscuously biotinylate proximal proteins by releasing an unstable intermediate^8, 9^. Biotinylated proteins can then be identified via mass spectrometry, providing a powerful workflow for an unbiased, high-resolution mapping of protein-protein interaction.

Importantly, proximity labeling captures these interactions in their biological context. We thus hypothesized that if proximal protein labeling could be done in a Ca^2+^-dependent manner—with a labeling enzyme activated by elevated local Ca^2+^ levels—it would enable proteomic investigation of Ca^2+^ signaling microdomains.

In this study, we engineered a biotin ligase that switches conformation between inactive and active states depending on its Ca^2+^-binding status. This enzyme, named Cal-ID, biotinylates nearby proteins when local Ca^2+^ levels are elevated. We expressed Cal-ID in HEK293T cells and investigated biotinylated proteins, allowing us to identify CEP131 and ASPM as primary components of the centrosomal Ca^2+^ signaling microdomain during mitosis. The localization of Ca^2+^ signaling microdomains and Cal-ID labeling exceeds the resolution of light microscopy, so we employed expansion microscopy technology to visualize ER Ca^2+^ release hotspots on the ER membrane. We also applied Cal-ID to mouse primary cortical neurons and found that plasma membrane Ca^2+^ ATPases (PMCA) 1 and 2 occupy the most prominent Ca^2+^ signaling microdomains. Cal-ID expressed in the mouse brain allowed us to distinguish activated neurons during an initial phase of kainic acid-induced seizure. These results highlight the value of Cal-ID as a novel and powerful tool that provides a new way to trace molecular events associated with local Ca^2+^ concentration in cells, including Ca^2+^ influx associated with neuronal activity.

## Results

### Development of Cal-ID, a Ca^2+^-dependent, promiscuous biotinylation ligase

The genetically encoded Ca^2+^ indicator GCaMP functions via a Ca^2+^-dependent conformational change that allows reversible intramolecular protein complementation of split green fluorescent protein (GFP). We envisioned that Ca^2+^-dependent complementation of a split proximity labeling enzyme, instead of split GFP, would enable proximity protein labeling in response to elevated local Ca^2+^ levels. We chose the BioID labeling enzyme, which is an R118G mutant of the BirA biotin ligase, because it does not require any toxic or synthetic substrates and is proven to work in living mouse brain^10–12^. Based on the configuration of the genetically encoded calcium indicator GCaMP6s, we fused the Ca^2+^ sensor calmodulin and the Ca^2+^- dependent calmodulin-binding peptide RS20 with circularly permuted, split BioID^13–15^ (Extended Data Fig. 1a). We expressed these fusions in HEK293T cells and found that a variant split between T195 and G196 has increased levels of promiscuous biotinylation when cytosolic Ca^2+^ levels were elevated by treatment with the sarco/endoplasmic reticulum Ca^2+^-ATPase (SERCA) inhibitor thapsigargin (Extended Data Figs. 1b and 1c). Of note, this T195/G196 split site has not been tested in previous studies of split BirA variants^14–17^. Since this engineered enzyme successfully responded to elevated cytosolic Ca^2+^ levels in living cells, we named it Cal-ID (Calcium-dependent BioID).

Because Cal-ID labeling is irreversible, background levels of Ca^2+^ -independent activity pose a more serious challenge than background signal in fluorescence-based Ca^2+^ indicators. Biotinylated proteins may accumulate over extended labeling periods, or in response to high Cal-ID expression levels. Therefore, we set out to suppress activity of Cal-ID in the low-Ca^2+^ state and to enhance its signal-to-noise ratio. First, since a highly active, promiscuous BirA derivative called TurboID has been developed, we introduced the TurboID mutations to Cal-ID^8^. Next, we designed an inhibitory peptide, AviTag*—a non-biotinylatable and truncated variant of the wild-type BirA substrate peptide AviTag—that should block the enzyme active site. We fused this pseudosubstrate peptide to the N-terminus of Cal-ID, where it inhibits the enzyme in the absence of Ca^2+^, yet is sequestered in response to Ca^2+^-dependent calmodulin-RS20 binding (Fig. 1a). Indeed, we found that the N-terminal AviTag* fusion increased the Ca^2+^ responsiveness of Cal-ID during an extended 1-hour incubation in high extracellular biotin (Extended Data Fig. 2a). Last, a recent study reported a new split site, L73/G74, for protein-fragment complementation of TurboID^16^. We revised Cal-ID with this split site since the new split site variant shows prominent biotinylation only after a 15-min incubation (Extended Data Fig. 2b).

**Fig. 1.**
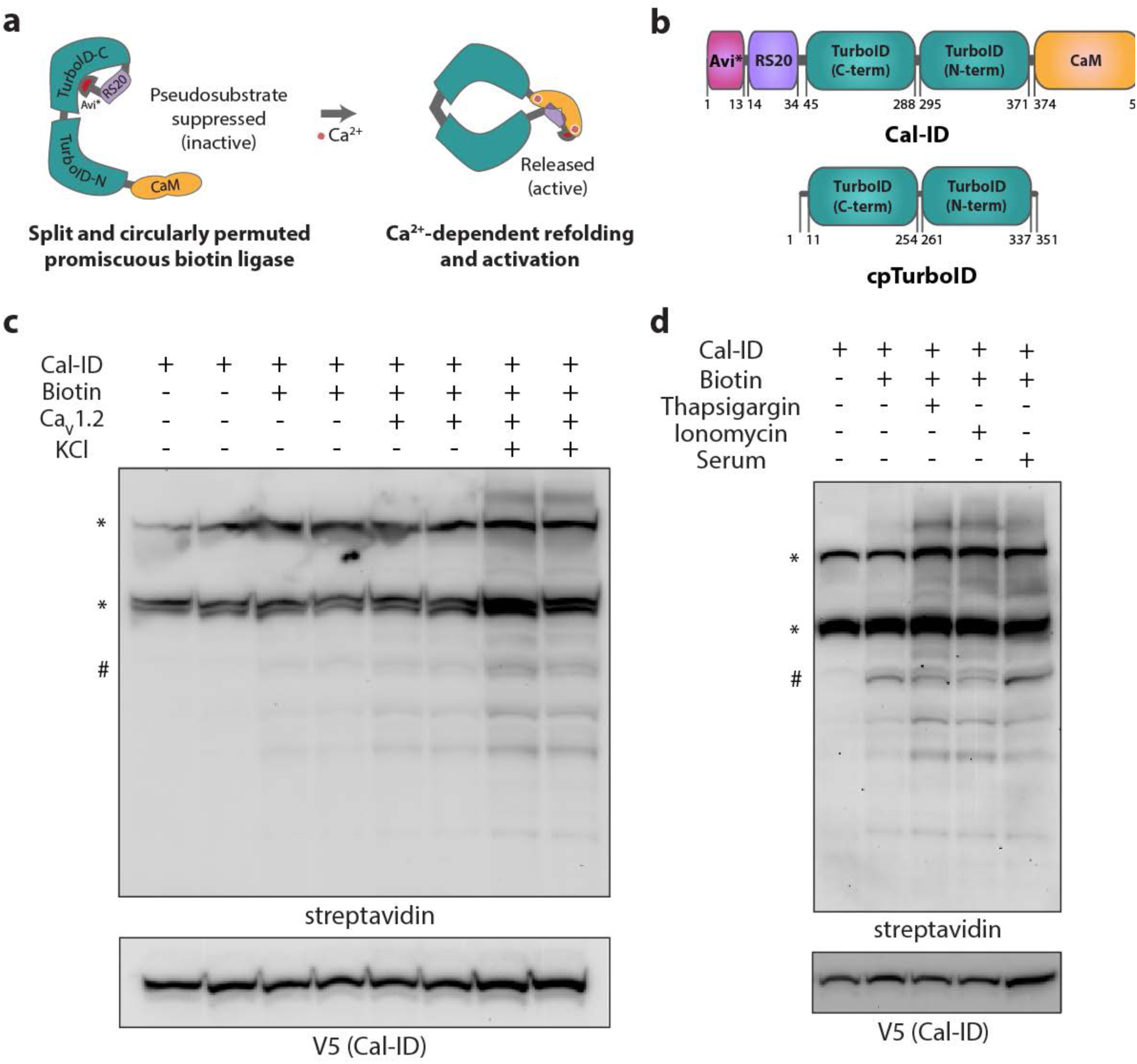
Cal-ID: a calcium-dependent proximal labeling enzyme. (**a**) A schematic diagram of Cal-ID activation. CaM: calmodulin, RS20: CaM-binding peptide. (**b**) Illustrations of Cal-ID and cpTurboID domain structure. Avi*: non-biotinylatable (K to R), truncated AviTag (NDIFEAQRIEWH). (**c** and **d**) Representative Cal-ID activation results in HEK293T cells. (c) Biotin: 50 µM, KCl: 80 mM. 30 min incubation. (d) Biotin: 50 µM, thapsigargin: 3 µM, ionomycin: 10 µM, serum: fresh media containing 10% FBS. 30 min incubation. *: endogenous biotin carrier proteins, #: self-biotinylation of Cal-ID.

We then tested Ca^2+^-dependent proximal biotinylation by this revised Cal-ID using various modes of cytosolic Ca^2+^ induction. In order to mimic the activity-dependent Ca^2+^ influx in neurons, we expressed the L-type voltage-gated calcium channel (L-VGCC) pore-forming alpha 1C subunit (CaV1.2) in HEK293T cells and showed that KCl depolarization triggers Cal-ID biotinylation (Fig. 1c). We also showed that Ca^2+^ influx mediated by the Ca^2+^ ionophore ionomycin also activates Cal-ID, similar to the effects of thapsigargin (Fig. 1d). Beyond these artificial stimuli, we observed that replenishing serum in the cell culture media also activates Cal-ID (Fig. 1d), suggesting that the activation of signaling cascades, which ultimately trigger ER-mediated Ca^2+^ release, represents a physiological stimulus that activates Cal-ID.

### Cal-ID revealed cell cycle-dependent Ca^2+^ microdomains at centrosomes

Because TurboID has much higher activity than Cal-ID, it is not an ideal point of comparison for Cal-ID labeling experiments. We thus constructed cpTurboID, a circularly permuted TurboID without calmodulin or RS20, as a Ca^2+^-independent control enzyme with labeling activity more similar to that of Cal-ID (Fig. 1b). cpTurboID is activated by spontaneous refolding, thereby providing a better control for the background activity of Cal-ID. It is also expected to reduce the risks of cellular stress from depleting cellular biotin over extended incubation time, which occurs with highly reactive TurboID^8^. To compare these enzymes at defined expression levels, we generated stable HEK293T cell lines with tetracycline-inducible Cal-ID or cpTurboID expression via Sleeping Beauty transposition and performed biotinylation experiments^18, 19^. We found that cpTurboID has stronger yet comparable biotinylation levels to Cal-ID when they are expressed at similar levels and incubated for the same amount of time (Fig. 2a). In addition, we found that treatment of BAPTA-AM, a cell membrane-permeable form of the Ca^2+^ chelator BAPTA, reduces Cal-ID-mediated biotinylation but does not affect cpTurboID-mediated biotinylation in cells (Extended Data Fig. 2c). These results demonstrate that cpTurboID is a suitable Ca^2+^-independent control for Cal-ID.

**Fig. 2.**
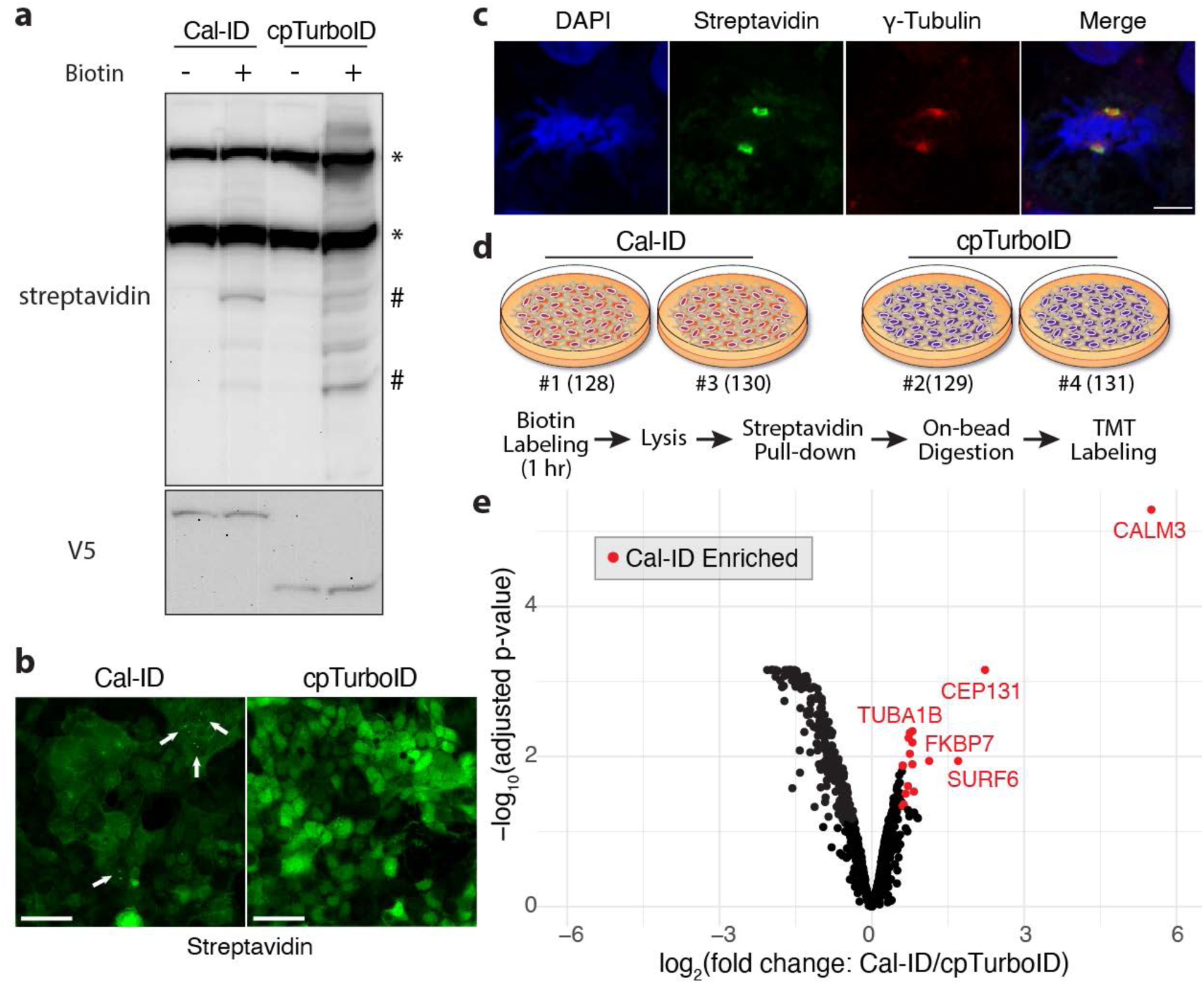
Identification of centrosomal calcium signaling microdomains proteins in mitosis. (**a**) Representative Cal-ID and cpTurboID biotinylation results from tet-inducible stable HEK293T cell lines. Induction: 400 ng/mL (Cal-ID), 200 ng/mL (cpTurboID) doxycycline for 48 hours. Biotin: 100 µM, 1 hour incubation. *: endogenous biotin carrier proteins, #: self-biotinylation of Cal-ID. (**b**) Streptavidin staining of Cal-ID- or cpTurboID-expressing stable HEK293T cell lines. Biotin: 100 µM, 1 hour incubation. Scale bar: 50 µm. (**c**) Centrosomal biotinylation of mitotic cells by Cal-ID. γ-tubulin is centrosomal marker. Biotin: 100 µM, 1 hour incubation. Scale bar: 5 µm. (**d**) A schematic diagram of TMT labeling for quantitative mass spectrometry. Isobaric tags: 128/130 (Cal-ID), 129/131 (cpTurboID). (**e**) A volcano plot showing Cal-ID enriched proteins discovered by mass spectrometry. Data points: 964. Differential expression between Cal-ID and cpTurboID was analyzed using limma with FDR 5% and log2 fold change > 0.5 (significant: red).

We next visualized the subcellular labeling pattern in Cal-ID-expressing cells using streptavidin-conjugated fluorophores and compared them with cpTurboID-expressing cells. cpTurboID biotinylation was observed throughout the cytosol, matching our expectations for a free and cytosolic promiscuous biotin ligase (Fig. 2b). In contrast, many Cal-ID-expressing cells showed distinctive, cell cycle-associated subcellular labeling patterns. In particular, mitotic Cal-ID-expressing cells showed strong biotinylation at the centrosome, which was visualized by γ- tubulin (Figs. 2b; see arrows, and 2b). Indeed, the importance of a Ca^2+^ microdomain in mitotic spindle formation is well established, and the elevated local Ca^2+^ concentration at the centrosome has been observed via genetically encoded Ca^2+^ indicators^20–23^. By detecting these well-studied Ca^2+^ microdomains, our results demonstrate that Cal-ID can detect physiologically relevant Ca^2+^ signaling events and demonstrate the spatial resolution of Ca^2+^-dependent, proximal protein labeling by Cal-ID.

Quantitative proteomics identified Ca^2+^ signaling constituents in living cells To investigate the distinct Cal-ID biotinylation patterns observed by microscopy in more detail, we wished to identify Cal-ID target proteins by proteomic mass spectrometry. Proximity labeling produces complex proteomic samples, but we and others have shown that ratiometric comparison of biotinylated samples can reveal specific patterns of biotinylation enrichment in the cell^24–26^. We thus carried out proximity labeling in Cal-ID- and cpTurboID-expressing cells and prepared samples for quantitative mass spectrometry with tandem mass tag (TMT) labeling for a ratiometric comparison of biotinylated proteins (Fig. 2d).

We detected 964 proteins overall, and biological replicates showed high quantitative reproducibility in both labeling enzymes (*r^2^* > 0.98; Extended Data Figs. 2d and 2e). We then compared cpTurboID- and Cal-ID-enriched proteins and identified 29 proteins that were significantly enriched in Cal-ID relative to cpTurboID samples at 5% false discovery rate (FDR) and log2 fold change > 0.5 cutoff (Fig. 2e, Supplementary Table 1). Cal-ID-enriched proteins are a subset of overall proteins labeled by cpTurboID, and so this analysis was designed to discover the most significant Cal-ID-specific labeling, but not to build an exhaustive list of the Ca^2+^- associated proteome. The dramatic enrichment of calmodulin presumably reflects Cal-ID self-biotinylation and recovery of tryptic fragments from this fusion protein. Due to its high sequence similarity, we cannot rule out the possibility that endogenous calmodulin is a major target of Cal-ID biotinylation, and so it could have contributed to the high enrichment of calmodulin as well. The second most enriched protein is CEP131, a centrosomal component protein. In addition to CEP131, ASPM, which is known to be associated with mitotic spindle regulation^27^, is also significantly enriched by Cal-ID labeling^27^. This result is consistent with the robust Cal-ID biotinylation at centrosomes in mitotic cells. In addition, we found that 5 out of 29 strong Cal-ID targets are tubulin ⍺- and β-subunits isoforms. Tubulins are the major component of the mitotic spindle, and β-Tubulin staining near the centrosome overlaps with fluorescent streptavidin staining (Extended Data Fig. 3a). Beyond CEP131, ASPM, and tubulins, Cal-ID enriches several known Ca^2+^ signaling-associated proteins. These labeled proteins include IQGAP1, a Ras GTPase-like protein with IQ domains that bind to calmodulin, and MCU, a mitochondrial Ca^2+^ transporter involved in mitochondrial Ca^2+^ uptake^28, 29^. We also saw enrichment of MYO6 and MYL6 myosin, which requires Ca^2+^ for their motor function. Last, we identified FKBP7, an ER chaperone that is thought to bind to Ca^2+^ because its mouse ortholog has Ca^2+^-binding affinity^30^.

Since several Cal-ID target proteins identified by mass spectrometry have known subcellular localization, we investigated their potential co-localization with Cal-ID biotinylation. We identified Cal-ID enrichment of two nucleolar proteins, SURF6 and ZCCHC17. We immunostained SURF6, which showed higher enrichment than ZCCHC17, and found that SURF6 was strictly localized to the nucleolus in non-mitotic cells, although streptavidin signal was relatively weak in the nucleolus (Extended Data Fig. 3b; top panel). However, during mitosis, SURF6 can be found in the cytosol and along the spindle fiber, suggesting that SURF6 biotinylation is mostly occurring during mitosis (Extended Data Fig. 3b; bottom panel). While the function of human SURF6 in mitosis is not clear, yeast orthologs of SURF6 are known to be involved in spindle formation, which potentially explains SURF6 localization and biotinylation during mitosis^31^. In addition, when we performed immunostaining of FKBP7, an ER protein, we were able to find a subset of cells that show ER biotinylation patterns overlapping with FKBP7 (Extended Data Fig. 3c). It is possible that these cells have active signaling cascades triggering Ca^2+^ release from the ER. In summary, these results collectively show that Cal-ID combined with ratiometric quantitative proteomics can identify physiologically relevant Ca^2+^ signaling microdomains in living cells.

### ER Ca^2+^ release hotspots visualized by expansion microscopy

Having characterized the strongest biotinylation targets of free cytosolic Cal-ID, we sought to test whether Cal-ID can be localized to an organelle in order to record local Ca^2+^ concentration. Particularly, we showed that serum replenishment activates Cal-ID potentially through ER Ca^2+^ release downstream of growth factor signaling (Fig. 1d). Thus, we hypothesized that Cal-ID targeted to the outer membrane of the ER would record Ca^2+^ concentration microdomains formed by ER Ca^2+^ release following serum replenishment. We generated ER outer membrane-targeted Cal-ID by adding the CYB5 ER transmembrane domain^32^ to the C-terminus of Cal-ID (Extended Data Fig. 3d). This enzyme, Cal-ID-ERM, showed robust activation by serum replenishment, which was completely blocked by BAPTA-AM treatment (Extended Data Fig. 3e).

We wished to visualize ER Ca^2+^ release by super-resolution microscopy as the size of Ca^2+^ microdomains can fall below the Abbe diffraction limit for light microscopy^2, 3^. We reasoned that expansion microscopy (ExM), which achieves effectively higher resolution by physically expanding a biological specimen^33^, could be readily applied to Cal-ID biotinylated samples because the biotinylated proteins and the streptavidin-fluorophore conjugates can be preserved through the physical expansion process. We labeled sites of serum-induced ER Ca^2+^ release in HEK293T cells using Cal-ID-ERM, expanded the cells by ∼4.5× using the protein retention ExM (proExM) method^34, 35^, and imaged the expanded sample with a confocal microscope (Fig. 3a). Imaging of biotinylated proteins using fluorophore-conjugated streptavidin visualized Ca^2+^ concentration ‘hotspots’ near the ER^36^ responding to serum activation, while the enzyme was expressed throughout the ER (Fig. 3b; top panel). Higher magnification images revealed that these hotspots are overlapping with Cal-ID-ERM epitope V5 signals or located adjacent to the V5 signals (Fig. 3b; bottom panel), suggesting that the streptavidin signals are coming from Cal-ID-ERM-biotinylated proteins along with the self-biotinylated enzyme. To examine how Ca^2+^ hotspots are positioned around ER Ca^2+^ release channels, we co-stained the expanded samples with antibodies against IP3 receptor, which is one of the main Ca^2+^ channels on the ER membrane for cytosolic Ca^2+^ release and involved in various Ca^2+^ signaling events including growth factor downstream signaling^36^. We visualized Ca^2+^ concentration hotspots around clusters of IP3 receptors (Figs. 3c and 3d, Supplementary Movies 1 and 2), demonstrating super-resolution visualization of biochemically recorded Ca^2+^ concentration microdomains near the ER. More broadly, we show that subcellularly targeted Cal-ID can generate a local record of Ca^2+^ concentration via biotinylation of proteins that can be visualized at super-resolution via ExM.

**Fig. 3.**
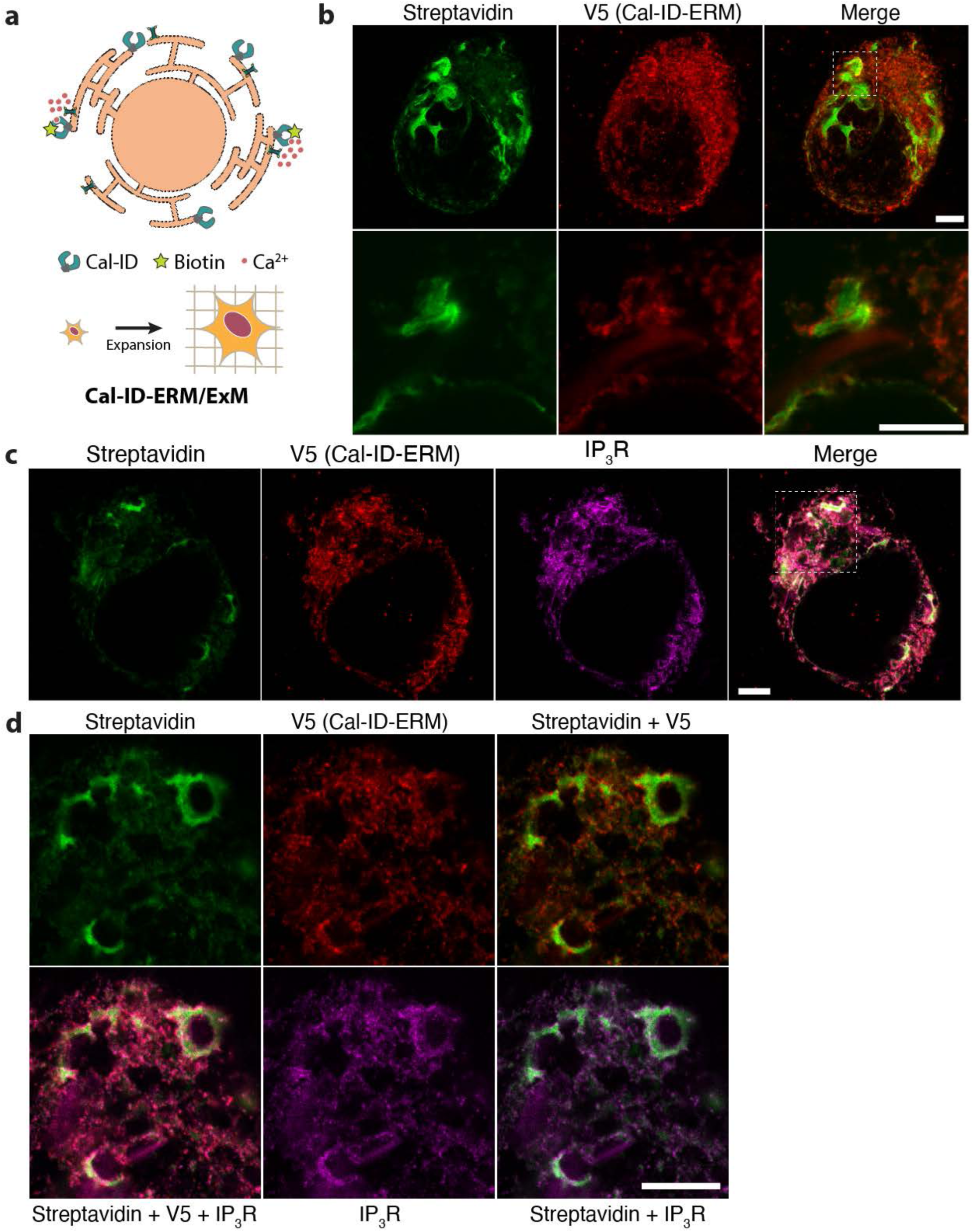
Visualization of ER calcium concentration hotspots using Cal-ID and expansion microscopy. (**a**) A schematic illustration of ER outer membrane-localized Cal-ID (Cal-ID-ERM) and expansion microscopy (ExM). (**b**) Representative images from Cal-ID-ERM-labeled and expanded HEK293T cells. Biotin: 100 µM, 20 min incubation post serum replenishment. Top panels: Z-projection of 7 images with 3 µm interval, bottom panel: single plane, both panels are from the same cell (see white box). (**c** and **d**) Representative images from Cal-ID-ERM-labeled and expanded HEK293T cells with V5/IP3R co-staining. Biotin: 100 µM, 20 min incubation post serum replenishment. (d) is a high magnification image from the same cell of (c) (see white box) but from a different plane. Cal-ID-ERM was transiently expressed. All scale bars: 2.2 µm (10 µm). For ExM images, the scale bar sizes are provided at the pre-expansion scale (the biological scale) with the corresponding post-expansion size (the physical size) indicated in parentheses.

### Cal-ID is activated by neuronal activity

In neurons, cytosolic Ca^2+^ influx from the extracellular space and release of Ca^2+^ from intracellular storage are key events in the regulation of neuronal function. Neuronal activity is tightly associated with Ca^2+^ influx through voltage-gated Ca^2+^ channels^37^. In addition, synaptic Ca^2+^ influx through neurotransmitter receptors, like NMDA receptors, is important for synaptic plasticity^37^. We therefore set out to express Cal-ID in mouse primary neurons and test whether Cal-ID biotinylation is increased by neuronal activity. We delivered Cal-ID or cpTurboID to mouse primary cortical neurons by lentiviral transduction (Fig. 4a). To test how Cal-ID responds to neuronal activity, we silenced or activated the neuronal cultures and compared the overall Cal-ID biotinylation levels via western blot. We silenced neuronal cultures with NBQX and APV, which are antagonists of AMPA and NMDA glutamate receptors, respectively. For activation, we first silenced the culture and then treated with CaCl2 to induce neuronal activity^38^. We found that the extent of Cal-ID mediated biotinylation tracked with neuronal activity in the culture, while cpTurboID-mediated biotinylation was unaffected by activity levels (Fig. 4b; see arrow).

**Fig. 4.**
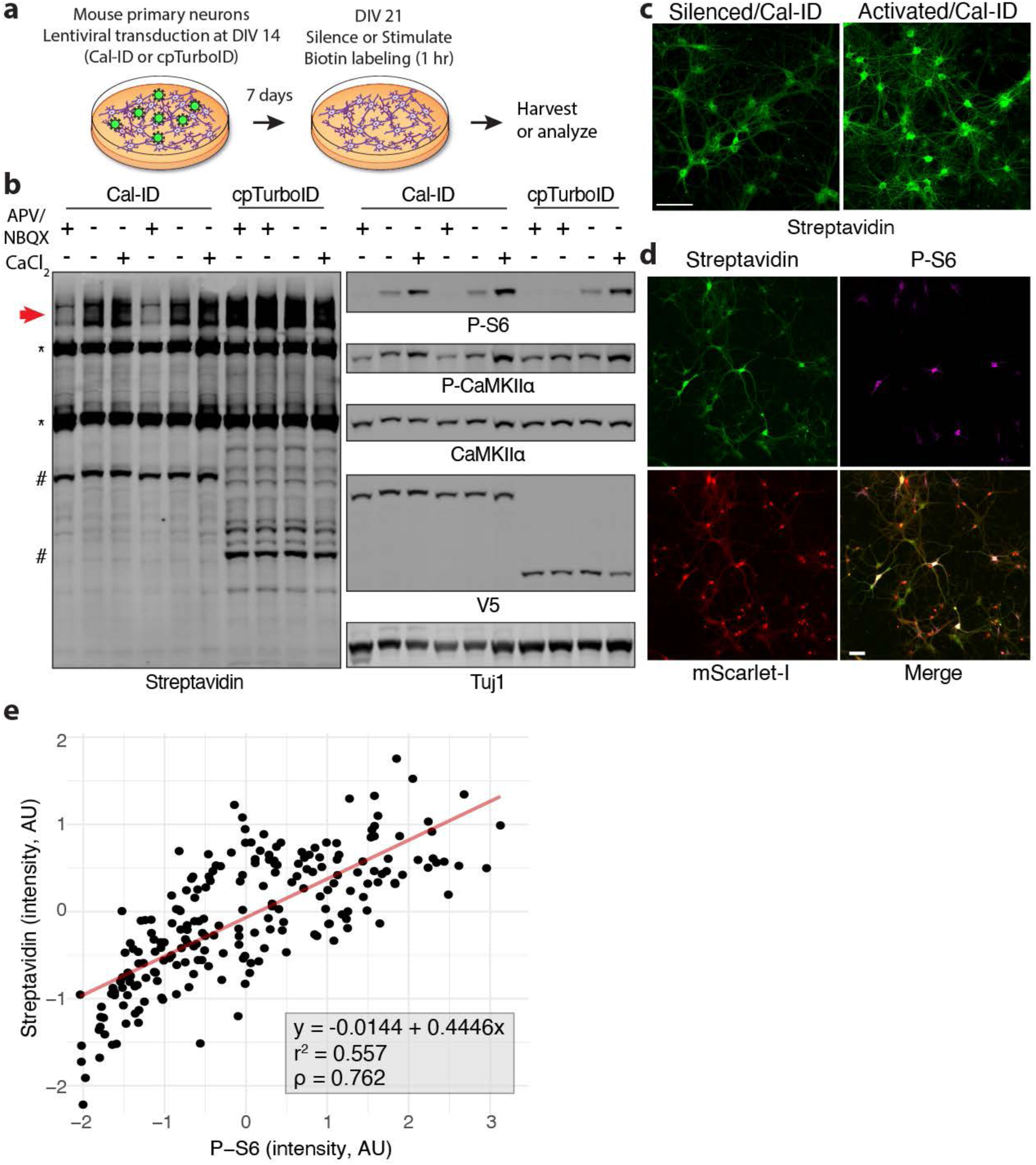
Cal-ID activation is associated with neuronal activity in neurons. (**a**) A schematic of Cal-ID or cpTurboID biotinylation in neurons. DIV: days in vitro. (**b**) Cal-ID biotinylation levels are associated with neuronal activity but cpTurboID biotinylation is not. 50 µM APV (NMDAR antagonist), 10 µM NBQX (AMPAR antagonist). APV + NBQX: treated overnight. 4 mM CaCl2: 1 hour. 100 µM biotin was added for 1 hour. P-S6 and P-CaMKII are activity markers. V5: Cal-ID or cpTurboID. *: endogenous biotin carrier proteins, #: self-biotinylation of Cal-ID or cpTurboID. (**c**) Comparison of overall biotinylation levels between silenced and activated conditions. Silenced: 50 µM APV + 10 µM NBQX for 2 hours, activated: 25 µM bicuculline (GABAAR antagonist) for 2 hours. 100 µM biotin for 1 hour. Mouse primary hippocampal neurons. Scale bars: 100 µm. (**d**) Representative images of P-S6 and streptavidin signals in mouse primary hippocampal neurons. mScarlet-I: Cal-ID expression control from Cal-ID-P2A-mScarlet-I construct. Scale bars: 50 µm. (**e**) Correlation of P-S6 and streptavidin signal intensity values from (d). Total 212 measurements from n = 3 (each condition). AU: arbitrary unit.

We next performed Cal-ID biotinylation in primary hippocampal neurons and visualized biotinylated proteins using streptavidin-conjugated fluorophores. We again found that activated neuronal cultures showed higher overall biotinylation levels than silenced cultures (Fig. 4c). In order to investigate cell-to-cell variation in Cal-ID labeling within a single culture and correlate it with neuronal activity, we generated a Cal-ID-P2A-mScarlet-I fusion that synthesizes mScarlet-I co-translationally with Cal-ID and splits the two proteins into separate polypeptides by means of the P2A sequence. After Cal-ID labeling, we stained neurons to visualize a classic marker of neuronal activity, phospho-rpS6 (Ser235/236), along with biotinylation^39^. We found that mouse primary hippocampal neurons expressing Cal-ID showed a strong positive correlation between streptavidin and phospho-rpS6 signal intensity (Figs. 4d and 4e). The expression levels of the Cal-ID protein, monitored by mScarlet-I, did not show a similar correlation with phospho-rpS6, so the difference in biotinylation reflects greater Cal-ID activity in neurons with high phospho-rpS6 rather than greater abundance of the enzyme (Extended Data Fig. 4a). Similarly, we also found a positive correlation between phospho-CaMKII and streptavidin signals but not from phospho-CaMKII and mScarlet-I signals from mouse hippocampal neurons (Extended Data Figs. 4b and 4c). The results indicate that Cal-ID labeling provides a biochemical strategy to study molecular events associated with neuronal activity that triggers intracellular Ca^2+^ influx.

### Plasma membrane Ca^2+^ ATPases are the strongest targets of Cal-ID in mouse primary neurons

We showed that Cal-ID labeled proteins in HEK293T cells have distinctive subcellular biotinylation patterns (Fig. 2b) and checked whether the same held true in neurons. We first performed subcellular fractionation from mouse cortical neurons to ensure that Cal-ID and cpTurboID have comparable subcellular localization patterns in the neuron. Fractionation showed that both Cal-ID and cpTurboID localize and biotinylate proximal proteins at all major target regions including cytosolic, membranous, presynaptic, and postsynaptic regions (Extended Data Fig. 5a). We then expressed Cal-ID or cpTurboID in primary hippocampal neurons and compared their biotinylation patterns using fluorescence microscopy. While both enzymes showed biotinylation across all components of the neuron, including the soma and the dendrites, fluorescent streptavidin staining revealed that Cal-ID-expressing cultures had more varying signal intensity levels between individual neurons compared to cpTurboID-expressing controls (Extended Data Fig. 5b; top panel). In addition, active neurons labeled by Cal-ID have a strong, saturating level of signals that centered at the soma and often include primary branches, while the overall signal intensity is comparable to that seen in cpTurboID-labeled neurons (Extended Data Fig. 5b, bottom panel).

We next sought to identify Cal-ID enriched proteins through a ratiometric proteomic comparison with cpTurboID. As in the experiment conducted with HEK293T cells, we prepared samples from mouse primary cortical neurons for quantitative mass spectrometry via TMT labeling. We captured 669 proteins in this labeling experiment, with high reproducibility (*r^2^* > 0.92) between biological replicates in both samples (Extended Data Figs. 6a and 6b). Similar to our analysis of the HEK293T mass spectrometry results, we again directly compared cpTurboID and Cal-ID enriched proteins to identify top targets of cytosolic Cal-ID in neurons and identified 6 proteins which were robustly biotinylated by Cal-ID with a 5% FDR (Fig. 5a). Calmodulin is again the protein most enriched by Cal-ID labeling, reflecting Cal-ID self-biotinylation and potential biotinylation of endogenous calmodulin as discussed above. We found two plasma membrane Ca^2+^ ATPases (PMCA1 and PMCA2; *Atp2b1* and *Atp2b2*, respectively) from the list of 6 significantly enriched proteins. PMCAs play a role in maintaining Ca^2+^ homeostasis by pumping Ca^2+^ out from the cytoplasm^40, 41^. The C-terminus of PMCAs is known to have multiple Ca^2+^ signaling-associated domains including calmodulin binding, high affinity Ca^2+^ binding for allosteric regulation, and a protein kinase C (PKC) target site^42, 43^. In addition to PMCAs, we identified wolframin (*Wfs1*), an ER membrane protein known to be involved in Ca^2+^ homeostasis, as a target of Cal-ID. Mutations in *Wfs1* cause Wolfram syndrome, which is characterized by juvenile-onset insulin-dependent diabetes mellitus with optic atrophy, deafness, dementia, and psychiatric illness^44, 45^. While the exact molecular mechanisms are unclear, it is thought to regulate ER Ca^2+^ levels in the context of ER-mitochondria crosstalk^46, 47^. These top Cal-ID targets identified by mass spectrometry demonstrated the capacity of Cal-ID to detect Ca^2+^ signaling microdomain-associated proteins in neurons.

**Fig. 5.**
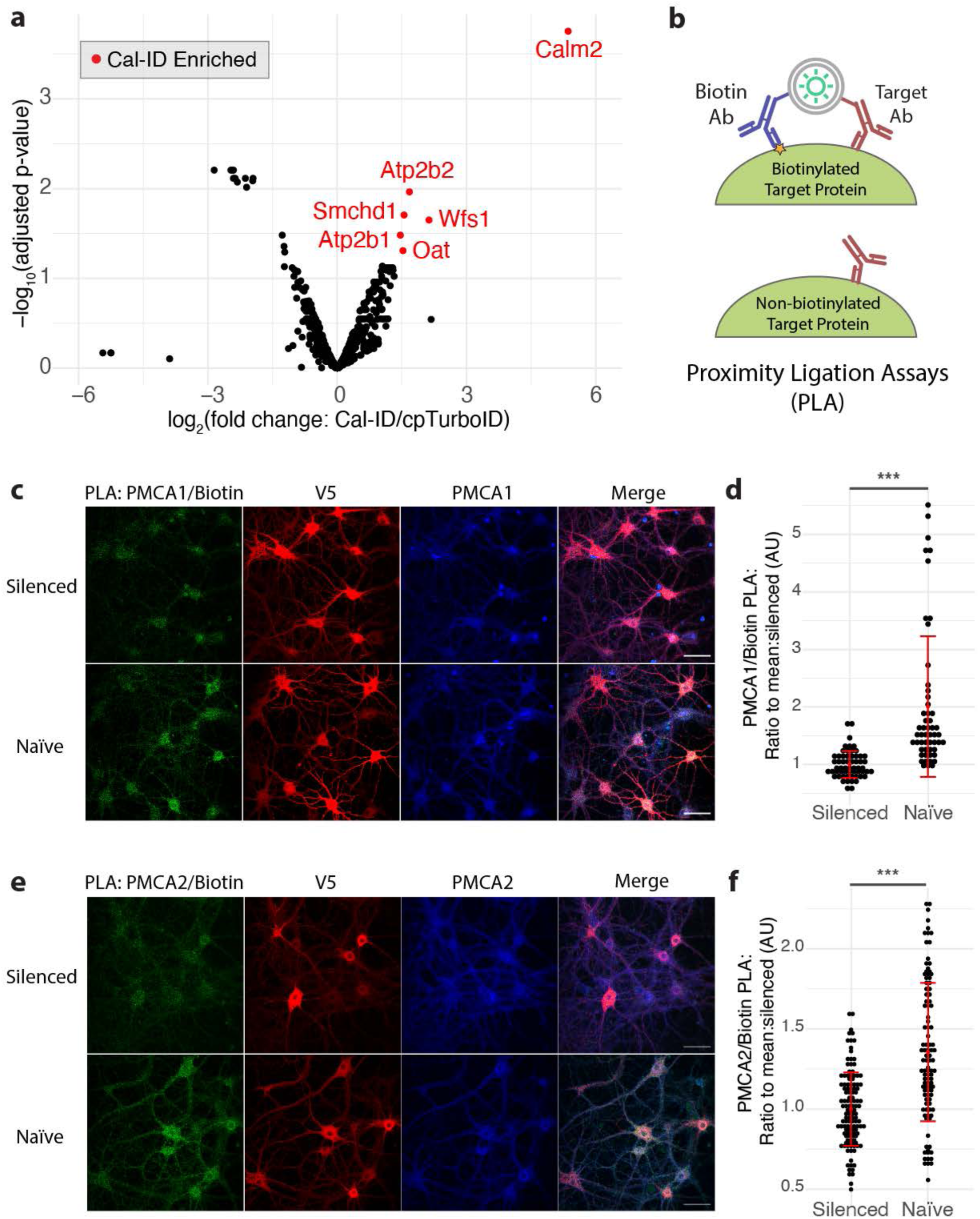
PMCA1/2 are associated with calcium signaling microdomains in neurons. (**a**) A volcano plot showing Cal-ID enriched proteins in mouse primary cortical neurons. Differential representation of 669 proteins recovered from both Cal-ID- and cpTurboID-expressing neurons was analyzed using limma with FDR 5% (significant Cal-ID enriched hits: red). Isobaric tags: 126/130 (Cal-ID), 127/131 (cpTurboID). (**b**) A schematic diagram of the proximity ligation assay (PLA). (**c** to **f**), PMCA1/2 and Biotin PLA. Silenced: 50 µM APV + 10 µM NBQX for overnight. Mouse primary hippocampal neurons, 100 µM biotin for 1 hour. (c) representative images of PMCA1/Biotin PLA, (d) quantification of (c), total 58 (silenced) and 53 measurements (naïve), n = 4 (each condition), (e) representative images of PMCA2/Biotin PLA, (f) quantification of (e), total 124 (silenced) and 105 measurements (naïve), n = 4 (each condition). Intensity ratios were calculated by normalizing raw intensity to the mean of the silenced condition for each batch. Statistical test: Student’s t-test, two-tailed; error bar: mean ± SD. AU: arbitrary unit. *** p < 0.001. Scale bars: 50 µm.

### Proximity ligation assay provides selective imaging of Cal-ID targets

As PMCAs were identified as the top Cal-ID target proteins, we hypothesized that biotinylation of PMCAs may have contributed to the distinct streptavidin-fluorophore staining patterns in neurons expressing Cal-ID (Extended Data Fig. 4b). Therefore, we sought to perform immunostaining of PMCAs and check their co-localization with streptavidin signals. However, the broad plasma membrane localization and high abundance of the PMCAs, as well as the presence of endogenous biotin carrier proteins, posed a challenge to the interpretation of co-localization. We therefore turned to proximity ligation assays (PLA), which rely on very close (< 40 nm) co-localization of two different antibodies to create fluorescent signal (Fig. 5b)^48^. PLA between antibodies for biotin and the target protein provides a complementary method to confirm biotinylation of a specific target protein by Cal-ID and resolve its location more clearly.

To confirm the applicability of PLA to visualize biotinylation in neurons, we first examined self-biotinylation of Cal-ID and cpTurboID enzymes using anti-biotin antibodies along with anti-V5 antibodies against the V5 epitope tag on the biotin ligases. Consistent with the previous streptavidin imaging and western blot results, we found an even starker difference in biotinylation between naïve and stimulated neurons expressing Cal-ID, but not from cpTurboID-expressing neurons (Extended Data Figs. 6c and 6d). Next, we performed PLA with biotin and PMCA antibodies. We found that biotin-PLA signals from both PMCA1 and PMCA2 were significantly decreased when neuronal activity was suppressed (Figs. 5c to 5f). By focusing on biotin derived from proximity labeling and excluding endogenous biotin, these PLA results verify that Cal-ID biotinylation is increased by neuronal activity.

### Cal-ID enables transcription-independent *in vivo* neuronal activity mapping

Brain-wide activity mapping strategies typically rely on reporter expression that requires transcription^49^. Our imaging and mass spectrometry results from neuronal culture revealed that Cal-ID robustly biotinylates plasma membrane proteins in active neurons, suggesting that Cal-ID biotinylation can be used for transcription-independent *in vivo* activity mapping in the brain.

Biotin readily crosses the blood-brain barrier, and BirA-based proximal biotinylation in the brain has been reported with exogenous supplementation of biotin^12, 50–52^. However, we noted that previous studies performed multiple rounds of biotin administration over several days to obtain biotinylated proteins.

Because multi-day labeling would not provide the high temporal resolution needed for brain activity mapping, we first tested whether a single biotin injection can produce detectable levels of biotinylated proteins *in vivo*. We expressed the constitutive cpTurboID labeling enzyme in the mouse brain by injecting adeno-associated virus (AAV) encoding cpTurboID into the cortex of neonatal mice at postnatal day 0 or 1 (P0/1). At 4 weeks of age, we intraperitoneally (i.p.) injected biotin and allowed biotinylation to proceed for 1 to 3 hours before sacrificing animals. Robust protein biotinylation was detected from all timepoints, including 1 hour labeling (Extended Data Fig. 6e), indicating that biotinylation began within 1 hour of biotin i.p. injection.

We then tested whether Cal-ID labeling and biotin imaging can demarcate activated neurons. After delivering Cal-ID by intracortical, unilateral AAV injection of P0/1 pups and allowing animals to mature for six weeks, we tested broad neuronal activation in the kainic acid-induced seizure model^53^. We performed biotin injection (i.p.), waited for 45 minutes to ensure that exogenous biotin reached the cortex, and then injected saline or kainic acid (i.p.) for the control or seizure conditions, respectively. We then monitored the mice for 45 minutes after the saline/kainic acid injection, ensured seizure behaviors (modified Racine scale 1 or higher^54^) in the seizure condition, euthanized the animals, and prepared brain tissue for fluorescence microscopy (Fig. 6a). We found prominent neuronal biotinylation in the cortex of kainic acid-injected mice, but not from the cortex of saline-injected mice nor the hemisphere without AAV injection (Fig. 6b, Extended Data Figs. 7a and 7b), showing that Cal-ID neuronal biotinylation can effectively report on induced activity in the brain. We quantified the streptavidin signal intensity in Cal-ID-expressing neurons across 6 kainic acid-injected mice and found that it was significantly higher than the signal seen in Cal-ID-expressing neurons across 5 saline-injected mice (Figs. 6c and 6d). These *in vivo* results show that Cal-ID biotinylation offers a new way to map brain activity that is independent of transcription-based reporter expression and can be triggered by biotin injection.

**Fig. 6.**
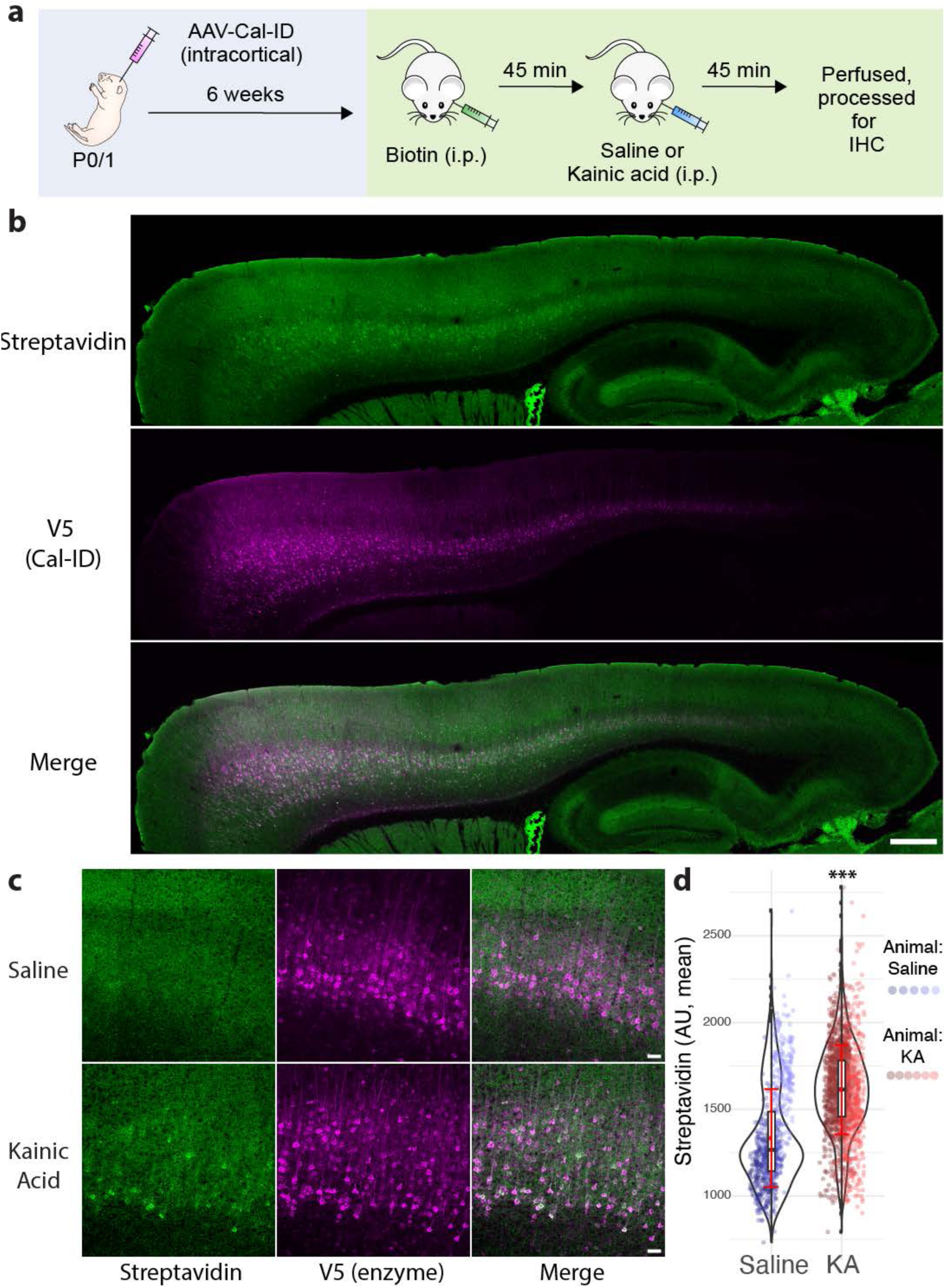
*In vivo* Cal-ID labeling to visualize active neurons in the brain. (**a**) A schematic of *in vivo* Cal-ID activity labeling of activated cortical neurons by kainic acid-induced seizure. AAV was injected to the cortex of postnatal day 0 or 1 CD1 mouse pups. i.p: intraperitoneal. Biotin: 24 mg/kg, kainic acid: 15 mg/kg. IHC: immunohistochemistry. (**b**) Representative images from a brain slice of kainic acid-injected and Cal-ID-labeled mouse brain. Sagittal section with 40 µm thickness. Scale bar: 500 µm. (**c**) Representative images from saline- or kainic-acid injected mouse brain slices. Scale bar: 50 µm. (**d**) Quantification of streptavidin mean signal intensity from Cal-ID expressing (V5 positive) neurons. Data points: 768 neurons from 5 mice (saline), 1874 neurons from 6 mice (kainic acid). AU: arbitrary unit. KA: kainic acid. Raw intensity values were obtained using the identical acquisition conditions. Statistical test: two-way ANOVA, [saline vs kainic acid] (F = 1062.1), error bar: mean ± SD. ***p < 0.001.

## Discussion

Ca^2+^ signaling is fundamental to cell physiology and widely conserved across eukaryotes. Most Ca^2+^ signals start from a transient increase of Ca^2+^ at a specific cellular region^2^. While Ca^2+^ diffuses rapidly, elevated local Ca^2+^ concentrations can be captured by Ca^2+^ sensor proteins and activate downstream signaling cascades. In this study, we exploited the function of the Ca^2+^ sensor protein calmodulin to control the activity of a promiscuous biotin ligase in a Ca^2+^-dependent manner. Since BirA-mediated biotinylation has a short labeling radius (∼10 nm), this strategy enables us to biochemically label proteins near the sites where local Ca^2+^ levels are elevated^7^ (Extended Data Fig. 8). Increased local Ca^2+^ occupies Ca^2+^ binding sites at calmodulin, shifting Cal-ID to the active, folded state via calmodulin-RS20 binding, thereby leading to proximal biotinylation. Once local Ca^2+^ levels drop to resting state, calmodulin will release Ca^2+^, disrupting the interaction with RS20 and allowing the Cal-ID to return to an inactive conformation. This conformational cycle places a permanent covalent mark on proximal proteins in response to transient changes in local Ca^2+^ concentration.

Cal-ID labeling in HEK293T cells highlighted Ca^2+^ signaling microdomains at the mitotic centrosome. The necessity of local Ca^2+^ signaling for mitotic spindle formation has been known for decades, which aligns well with our observation of robust Cal-ID biotinylation at the centrosome. It is noteworthy that, while multiple pericentriolar proteins such as pericentrin (PNCT) and AKAP450 contain potential calmodulin-binding motifs, the primary proteins constituting centrosomal Ca^2+^ signaling microdomains have not been identified^55, 56^. Our Cal-ID mass spectrometry results implicated CEP131 and ASPM—both of which contain a calmodulin-binding IQ domain—as key components for centrosomal Ca^2+^ signaling. Potential Ca^2+^- dependent roles of CEP131 and ASPM in mitotic spindle regulation could expand our knowledge of Ca^2+^ signaling events at the centrosome.

Ca^2+^ transients on the outer surface of the ER, which have been seen in many different cell types, often show ‘hotspots’ of Ca^2+^ release^2^. These hotspots are typically generated at ER-plasma membrane or ER-mitochondria junctions, and they are thought to be the sites of local Ca^2+^ signaling^36, 57^. In this study, we recorded ER Ca^2+^ release hotspots using Cal-ID-ERM and visualized them at super-resolution with ExM technology. With 20 minutes of recording, we were able to show strongly biotinylated proteins near the ER membrane and around some IP3R Ca^2+^ channels, and potential diffusion or transport of these proteins from the original biotinylated sites. We demonstrated that the combination of these two technologies synergizes powerfully; ExM enables visualization of subcellular Ca^2+^ microdomains recorded by Cal-ID at super-resolution with relatively high accessibility, as imaging can be performed with conventional diffraction-limited microscopes.

In neurons, Ca^2+^ influx is associated with neuronal activity and plays an essential role in many regulatory processes^4^. At synapses, Ca^2+^ influx via neurotransmitter receptors, Ca^2+^ channels, and release of ER-stored Ca^2+^ is a central signaling mechanism for local synaptic regulation on both short (milliseconds) and long (hours) timescales^58^. Ca^2+^ signaling back to the nucleus is essential for activity-dependent gene expression^59^. We therefore expect that identifying Ca^2+^ microdomain constituents via Cal-ID will advance our understanding of the molecular physiology of the neuron. In primary cortical neurons, Cal-ID labeling enriches Atp2b1 (PMCA1) and Atp2b2 (PMCA2)—major Ca^2+^ pumps which are responsible for extruding cytosolic Ca^2+^. The high abundance of PMCAs in neurons and their affinity to calmodulin may contribute to their Cal-ID labeling. However, there are other abundant neuronal proteins with known calmodulin-binding motifs, such as voltage-gated Ca^2+^ channels and ryanodine receptors, which drive major Ca^2+^ influx into cytoplasm in neurons^37^. Therefore, it is plausible to speculate that high local Ca^2+^ concentrations and Ca^2+^ signaling microdomains are maintained around PMCAs, since active export is relatively slower than Ca^2+^ influx through channels. In addition, our results helped to speculate the subcellular location of fluorescent signals generated by genetically encoded Ca^2+^ indicators that incorporate calmodulin, such as GCaMPs. Further investigation of the Ca^2+^ signaling events around PMCAs is warranted to expand our understanding on the formation and regulation of Ca^2+^ microdomains in neurons.

With prominent plasma membrane biotinylation that facilitates visualization of Cal-ID-labeled neurons, Cal-ID labeling and imaging could be especially advantageous in neuroscience as an approach to track neuronal activity. Cal-ID biotinylation can be temporally controlled through biotin delivery and reaches detectable levels within 1 hour of labeling. We showed that Cal-ID biotinylation in the mouse cortex was elevated during induced activity caused by kainic acid treatment, leading to prominent and non-uniform labeling of individual neurons. These *in vivo* results show that Cal-ID provides a new strategy for brain activity mapping by attaching covalent biochemical marks to nearby proteins in response to Ca^2+^ influx, instead of expressing reporter transcripts or proteins under control of the Ca^2+^ signaling cascade. The nature of proximity labeling gives Cal-ID the potential to record neuronal activity at a subcellular level—perhaps even at individual synapses—across the whole brain. Furthermore, proteomic analysis of biotinylated proteins will enable identification of proteins associated with local neuronal activity in the brain. Of note, this approach may offer distinctive advantages over other tools such as CaMPARI, a photoactivatable Ca^2+^ indicator^60, 61^. While CaMPARI can ‘mark’ neuronal activity at a great temporal resolution, the need for light activation limits the maximum area of recording. However, enzymatic labeling by Cal-ID will be much slower than the millisecond timescale of CaMPARI, which must be considered when interpreting these data.

More broadly, Cal-ID differs in important ways from fluorescent Ca^2+^ indicators. Cal-ID functions on a different timescale and may require incubation over many minutes to accumulate detectable signals. Therefore, while Cal-ID is activated by increased Ca^2+^ concentration, Cal-ID biotinylation does not necessarily represent a single Ca^2+^ transient. When biotinylation levels are low, visualization of Cal-ID biotinylation may benefit from the amplification and specificity offered by PLA or other approaches, because endogenous biotin carrier proteins give background biotin signals from mitochondria. Additionally, it has been reported that ectopic expression of Ca^2+^-binding proteins in fluorescent indicators can alter Ca^2+^ buffering, and Cal-ID may have similar limitations^62^. Finally, due to the central roles of calmodulin in Ca^2+^ signaling, it is possible that Cal-ID localization is affected by the inclusion of calmodulin, and proteins with calmodulin-binding motifs are more easily targeted for biotinylation. Since many calmodulin-binding partners have Ca^2+^-dependent binding affinity, and most calmodulin-associated molecular processes are involved in Ca^2+^-dependent regulation, we believe that the possibility of identifying false positives due to calmodulin-binding affinity is low^63^. Nonetheless, we suggest that Cal-ID labeling results should be interpreted with due consideration for the existence of calmodulin-binding motifs.

In this study, we presented Cal-ID, an engineered proximal protein ligase that is activated by Ca^2+^-dependent refolding. Cal-ID functions as a molecular recorder of Ca^2+^ microdomains, thereby providing a unique way to investigate Ca^2+^ signaling events in cells. We showed that Cal-ID-labeled proteins can be analyzed in many different ways by employing technologies such as mass spectrometry, proximity ligation assay, and expansion microscopy. We also demonstrated the *in vivo* capability of Cal-ID by successfully labeling neurons in the activated mouse brain. Thus, we suggest that Cal-ID has great application potential in multiple fields of biology in which cellular activation accompanied by Ca^2+^ influx is important.

## Methods

All animal protocols are in accordance with the regulations of University of California, Berkeley Animal Care and Use Committee and the National Institutes of Health (NIH) *Guide for the Care and Use of Laboratory Animals.* Animals were housed in a 12-hour reverse dark/light cycle with free access to water and food.

### Cloning

For cloning, PCR primers were designed to have overhangs, PCR reactions were performed with Q5 polymerase (New England BioLabs, NEB), and the fragments were assembled using HiFi (NEB) or Gibson assembly (NEB). Ligated plasmid products were introduced by heat shock transformation into competent Stbl3 (Invitrogen). The cloning products were confirmed using Sanger sequencing. Calmodulin and RS20 sequences were derived from GCaMP6s^13^. All biotin ligase variants were derived from E. coli biotin protein ligase, from BioID or TurboID^8, 9^.

### HEK293T Cell Culture and Generation of Stable Cell Lines

HEK293T cells were maintained with DMEM with GlutaMax (Gibco) supplemented with 10% tetracycline-free FBS (Gibco). Cal-ID and cpTurboID transgenes were cloned into the pSBtet-Hyg (Addgene #60508) donor vector, mixed at a 5:1 ratio with pCMV(CAT)T7-SB100 (Addgene #34879), and co-transfected to HEK293T cells using TransIT-293 (Mirus). 48 hours after transfection, 200 μg/mL hygromycin (Invitrogen) was added to the culture medium and selected for 3 passages each with a 1:20 split ratio.

### Primary Neuron Culture

Dissociated primary cortical neurons were prepared from E15 developing brain (CD1, Charles River). Developing cortices were dissected in dissecting medium (Dulbecco’s Modified Eagle Medium (DMEM) with 20 % FBS, 0.5 mM GlutaMax, 6 μM glucose, Gibco), digested with 20 mg/ml papain (Worthington) for 20 min at 37 °C, and plated at a concentration of 1.2 × 10^6^ cells for a 12-well plate (cortical neurons) or 60,000 – 70,000 neurons per one well of 24-well plate (hippocampal neurons), 4.2 × 10^6^ cells per 100 mm dish, 7 × 10^6^ cells per 150 mm dish. Culture plates were pre-coated with 50 μg/mL poly-D-lysine. Cultures were maintained under Neurobasal Plus (Gibco) medium with a serum-free supplement B-27 plus (Gibco) and 0.5mM GlutaMax (Gibco). All animal procedures were conducted according to protocols approved by the Institutional Animal Care and Use Committee (IACUC) and the Office of Laboratory Animal Care (OLAC) at the University of California, Berkeley (AUP-2018-08-11380).

### Immunocytochemistry

Cells were washed 3 times and fixed with 4 % paraformaldehyde for 15 min at RT, washed 3 times again then permeabilized and blocked for 1 hour with 5 % BSA and 0.1 % saponin in PBS. The blocked cells were subsequently incubated with primary antibody overnight at 4 °C. On the following day, the cells were washed 3 times and incubated with secondary antibody for 1 hour at room temperature in a light controlled condition. After 3 × wash with PBS buffer, the cells were mounted on cover slides with mounting media containing DAPI (VECTASHIELD® Plus, Vector Labs, H-1900). HEK293T and neuronal images were taken with LSM710 (Carl Zeiss) confocal laser scanning microscope under 20 × air or 63 × oil objectives or SP8 (Leica) confocal laser scanning microscope with a 40 × oil objective.

### Expansion microscopy

The pro-ExM procedure was performed following a previously published protocol^64^. Briefly, HEK293T cells on coverslips were labeled, fixed, permeabilized and stained as described in the immunocytochemistry method. Stained cells were treated with 0.1 mg/mL AcX overnight in the dark at RT. The following day, cells were washed with 1× PBS for 3 times. Then the coverslip was reverse mounted (cells facing downwards) on the cover glass gelation station with #1 thickness coverslip spacers and 10 mm opening in the middle, which is placed on ice; gelation solution was added to the coverslip and also the gelation station to fill out the space. Gelation solution is 98:1:1 mixture of monomer solution (1× PBS, 2 M NaCl, 8.6% (w/v) sodium acrylate (Pfaltz&Bauer), 2.5 % (w/v) acrylamide, 0.15 % (w/v) N,N’-Methylenebisacrylamide), tetramethylethylenediamine (TEMED) 10% (w/v) stock solution, and ammonium persulfate (APS) 10 % (w/v) stock solution. Gelation reaction was performed for 1 hour in a humidified 37 °C chamber. Sample gels were carefully removed from coverslips using a razor blade, transferred to a small petri dish using a fine paint brush, then immersed in Proteinase K digestion solution (0.5 % (w/v) Triton X-100, 1 mM EDTA, 50 mM Tris pH 8, 1 M NaCl, 8 U/mL Proteinase K (NEB)) and incubated overnight at RT. The following day, the digested gel samples were transferred to a larger petri dish then washed/expanded with sufficient ddH20 for 3 times, 20 minutes for each wash. Expanded gel samples were mounted on poly lysine-coated glass coverslips for imaging. Unless otherwise stated, all chemicals for ExM were from Millipore Sigma.

### Immunoblotting

Brain tissues were lysed with an automated homogenizer in RIPA buffer with 1 % SDS (20 mM Tris-HCl (pH 7.4), 150 mM NaCl, 1 mM EDTA, 1 % NP-40, 1 % sodium deoxycholate, 1 % SDS, protease inhibitors). Lysates were incubated on a rotator for 1 hour at 4 °C, and debris was separated by centrifugation for 10 min × 12,000 g at 4 °C. Supernatant was collected, protein concentration was measured, and the lysate was mixed with 2 × Laemmli sample buffer. SDS-PAGE and transfer were performed on Invitrogen Bolt™ system with Bis-Tris 4-12% gradient gels. ProteinSimple or Li-Cor imager was used for visualization.

### Lentivirus generation

Cal-ID or cpTurboID transgenes were cloned into pJW1511 (Addgene #62365) or FUGW (Addgene #14883) transfer vectors after removing the original transgene. Transfer vectors were co-transfected into low passage number (< 15) HEK293T cells with 3^rd^ generation VSV-G and gag/pol plasmids at a 1:2:3 ratio. 1 day after transfection culture, media was replaced with new media with 10 mM (final) HEPES. The first harvest of packaged virus was made 2 days after transfection, mixed with Lenti-X concentrator (Takara), and stored at 4 °C. A second harvest was made 3 days after transfection, mixed with Lenti-X concentrator and concentrated along with the first harvest following the manufacturer’s recommendation. The relative titer of the lentiviral concentrate was estimated using Lenti-X GoStix (Takara).

### Sample preparation for Mass Spectrometry

Samples for mass spectrometry were prepared following the previously published protocol for proteomics analysis of TurboID-labled proteins from Alice Ting’s laboratory^65^. Briefly, 2 × 150 mm dishes of cells were used for each sample. Biological replicates were prepared for both cpTurboID and Cal-ID labeling. For inducible HEK293T cell lines, 400 ng/mL doxycycline was used for Cal-ID induction and 200 ng/mL doxycycline was used for cpTurboID induction, each for 48 hours. For lentiviral transduction, concentrated lentiviral titer was estimated by Takara GoStix and GV ∼25,000 (cpTurboID) or ∼50,000 (Cal-ID) viruses were applied to neuronal cultures at 1:50 volume ratio. For biotinylation, 100 µM biotin was added and incubated for 1 hour in tissue culture incubator. Cells were washed with ice-cold PBS with 5 times and lysed with 1.5 mL RIPA lysis buffer (50 mM Tris, 150 mM NaCl, 0.1 % (wt/vol) SDS, 0.5 % (wt/vol) sodium deoxycholate, 1 % (vol/vol) Triton X-100, pH 7.4). 2.5 % of lysates were saved and further analyze by western blot to ensure biotinylation of the samples. The samples were incubated with 250 μl magnetic streptavidin beads (Thermo Fisher Scientific) at 4 °C overnight with rotation. The beads were washed twice with RIPA buffer (1 mL, 2 min at RT), once with 1 M KCl (1 mL, 2 min at RT), once with 0.1 M Na2CO3 (1 mL, ∼10 s), once with 2 M urea in 10 mM Tris-HCl (pH 8.0) (1 mL, ∼10 s), and twice with RIPA buffer (1 mL per wash, 2 min at RT). 2.5% of the beads were analyzed by silver staining to ensure successful pulldown of biotinytlated proteins. The beads were washed again with 200 μL of 50 mM Tris-HCl (pH 7.4), twice with 200 μL 2M urea in 50 mM Tris (pH 7.4) buffer. Then the beads were incubated with 80 μL of 2 M urea in 50 mM Tris-HCL containing 1 mM DTT and 0.4 μg trypsin (Promega) at 25 °C for 1 hour while shaking at 1,000 rpm. The supernatant containing digested peptides was saved, and beads were washed twice with 60 μL of 2 M urea in 50 mM Tris (pH 7.4) buffer and combined the washes with the digestion supernatant. DTT was added to a final concentration of 4 mM and incubated with the eluate at 25 °C for 30 min with shaking at 1,000 rpm. The samples were alkylated by iodoacetamide to a final concentration of 10 mM and incubated at 25 °C for 45 min with 1,000 rpm agitation, protected from light. 0.5 μg of trypsin was added to the samples and further digested overnight at 25 °C with shaking at 700 rpm. The next day, formic acid was added at ∼1 % (vol/vol) to adjust sample pH to ∼ 3. Samples were desalted using C18 StageTips as previously described^66^.

### TMT labeling and quantitative mass spectrometry

Peptides were quantified and normalized using a quantitative colorimetric peptide assay (Thermo Fisher Scientific). TMT reagents were reconstituted with 41 μL of anhydrous acetonitrile with occasional vortexing for 5 min at RT. The reduced and alkylated proteins were transferred to the TMT vials and incubated for 1 hour at RT. The reaction was quenched with addition of 8 μL of 5% hydroxyamine and incubation for 15 min. The samples were desalted with C18 StageTips. Samples were run on an OrbiTrap mass spectrometer (ThermoFisher Scientific) using high pH fractionation. Raw data from proteomics experiments were deposited in ProteomeXchange with identifier PXD033244.

### Mass Spectrometry Data Analysis

Mass spectra were processed using Proteome Discoverer (Thermo Scientific). Quantitative mass spectrometry results were analyzed using limma (3.34.9) in R^67^. Proteins with no peptide detected from any condition were omitted. Pre-normalization raw intensity values were transformed by the voom function, and the values were fitted for linear models via limma to estimate differential representation of proteins in Cal-ID vs cpTurboID pull-down samples with biological replicates. FDR was calculated by Benjamini-Hochberg correction.

### Proximity Ligation Assay

Proximity Ligation Assay (PLA) was performed using Duolink® (MilliporeSigma, DUO92101) following the manufacturer’s protocol. Briefly, mouse hippocampal neurons were fixed with 4 % paraformaldehyde for 15 minutes at RT, then permeabilized with 0.1% Triton X-100 for 15 min. The cells were washed then blocked for 1 hour at 37 °C with a humidity chamber with the Duolink® Blocking Solution. The blocked cells were subsequently incubated with primary antibodies diluted with the Duolink® Antibody Diluent (conditions are below) overnight at 4°C. The following day, cells were washed twice with Wash A buffer for 5 min at RT. Duolink® PLUS and MINUS probes were diluted with the Diluent with 1:5 concentration, added to the coverslips, incubated at 37 °C for 1 hour in a humidity chamber. After incubation, the coverslips were washed twice with Wash A buffer for 5 min at RT. When any additional secondary antibodies were used in addition to the PLA probes, secondary antibodies in the Diluent were added and incubated for 40 mins at 37 °C, and washed with Wash A buffer twice for 5 min.

After that, 1 × Duolink® Ligation buffer was prepared, ligase was added following 1:40 ratio, mixed well and added to the coverslips. The ligation reaction was performed for 30 min at 37 °C in a humidity chamber. Following ligation, coverslips were washed twice with Wash A buffer, and 1 × Amplification reaction mix with 1:80 enzyme:buffer ratio was prepared and added. The amplification reaction was performed for 100 min at 37 °C in a humidity chamber. Final wash was performed with 1 × Wash B buffer twice for 10 min then 0.01 × Wash B buffer once for 1 min. The coverslips were mounted using VECTASHIELD PLUS® (Vector Labs) or Duolink® DAPI mounting solution provided with the kit.

### Image Analysis

For streptavidin and P-S6/P-CaMKII correlation, PLA, brain slice analyses, ImageJ (1.53f51)/FIJI was used^68^. Cell bodies of the mScarlet-I or V5-expressing neurons were selected via Threshold, Watershed, and Analyze Particles tools with 50 – infinity size selection in ImageJ. Cells on the edges were excluded. After selecting soma regions, target mean intensity values were extracted from the selections and transferred to R. The results were analyzed in R and visualized using ggplot2 (3.0.0)^69^. The heatmap was generated by Calibration Bar tool in ImageJ. Movie was generated using IMARIS (Oxford Instruments, version 10).

### Subcellular Fractionation

Samples were kept on ice during the procedure and centrifugations and incubations were conducted at 4 °C. All solutions contained phosphatase inhibitors (Sigma-Aldrich PhosSTOP^TM^), protease inhibitors (Roche cOmplete^TM^), 5 mM sodium pyrophosphate, 1 mM EDTA, and 1 mM EGTA. Cultured cortical neurons were collected by scraping and homogenizing by passage through a 26G needle 12 times, in homogenization buffer (320 mM sucrose, 10 mM HEPES).

The homogenate was centrifuged at 800 g for 10 min to obtain the post-nuclear pelleted fraction 1 (P1) and supernatant fraction 1 (S1). S1 was further centrifuged at 15,000 g for 20 min to yield P2 (membranous fraction) and S2 (cytosolic fraction). P2 was resuspended in Milli- Q® water, adjusted to 4 mM HEPES (pH 7.4), and incubated with agitation for 30 min before centrifugation at 25,000 g for 20 min to obtain LP1 (mitochondria, pre- and postsynaptic membranes) and LS2 (synaptosomal cytosolic fraction). LP1 was resuspended in 50 mM HEPES (pH 7.4), mixed with an equal volume of 1 % Triton X-100, and incubated with agitation for 8 min. Lastly, centrifugation at 25,000 g for 20 min yielded the presynaptic membranes (supernatant) and the postsynaptic density (pellet).

### Intracranial neonatal mouse injections

CD1 neonatal mice (P0) were cryo-anesthetized by placing on ice for ∼2–3 min. When the animal was fully anesthetized, confirmed by lack of movement, it was gently placed in a head mold. Each pup received a total of 700 nL of 1:1 saline-diluted AAV9 (HHMI Viral Tools, Cal-ID: 7. 6×10^12^ GC/mL, cpTurboID: 5.1×10^12^) spread across 2 × 350-nL injections at a rate of 250 nL/minute. Two 350 nL unilateral injections were made into the left cortex, each one fourth the distance between bregma and lambda, approximately 0.6 mm from the sagittal suture and 0.4 mm ventral to the surface of the skull.

### Seizure induction

Male and female littermates were housed on a reversed light-dark cycle and six weeks after neonatal injections, were administered kainic acid or saline via intraperitoneal (i.p.) injection during the dark cycle. Prior to seizure induction/saline injection, all mice were first given 24 mg/kg of biotin diluted in sterile saline. Half of the mice were given then saline and the other half given 15 mg/kg kainic acid by i.p. injection to induce seizures 45 minutes after the initial biotin injection. Mice were monitored for 45 minutes post seizure induction/saline injection; seizure phenotypes were determined following modified Racine scale, higher than 1 (whisker trembling and/or facial twitching)^54^. Mice were perfused via transcardiac perfusion.

### Perfusion and immunohistochemistry

6 week old mice were deeply anesthetized by isoflurane and transcardially perfused with ice-cold 1× PBS, followed by 4 % PFA solution (Electron Microscopy Services: 15713) in 1× PBS using a peristaltic pump (Instech) 45 minutes post seizure induction/saline injection. The brains were removed and post-fixed by immersion in 4 % PFA in 1× PBS overnight at 4 °C. Brains were then suspended in 30 % sucrose in 1× PBS solution at 4 °C for approximately 24 hrs until the brains sunk. Brains were then sectioned sagittally at 40 µm on a freezing microtome (American Optical AO 860). Free-floating brain sections were batch processed to include matched saline and kainic acid injected samples. With gentle rocking on a nutator, sections were washed 3 times for 5 min in 1× PBS, followed by 1 hour incubation at RT with BlockAid blocking solution (Life Technologies B10710). Primary antibodies (V5: Invitrogen R960-25, 1:1000, Streptavidin-AF488: Invitrogen S11223, 1:750) were diluted in 1× PBS-T (1× PBS + 0.25% Triton-X-100) and applied overnight at 4 °C. Sections were washed with cold 1× PBS-T for 3 times each for 10 min then incubated for 2 hours at RT with secondary antibodies (AF546-Ms: Invitrogen A11003, 1:500) diluted in 1× PBS-T. Sections were then washed in 1× PBS 3 x 10 min, mounted on SuperFrost Plus slides (Fisher Scientific 12-550-15) and coverslipped with ProLong Gold antifade reagent with DAPI (Invitrogen P36935). Images of brain sections were taken on an FV3000 Olympus confocal microscope equipped with 405, 488, 561, and 640 nm lasers and a motorized stage for tile imaging. Z-stack images captured the entire thickness of the section at 1-2 µm steps using a 10 × air objective with a 2 × optical zoom.

**Supplementary Table 1.**
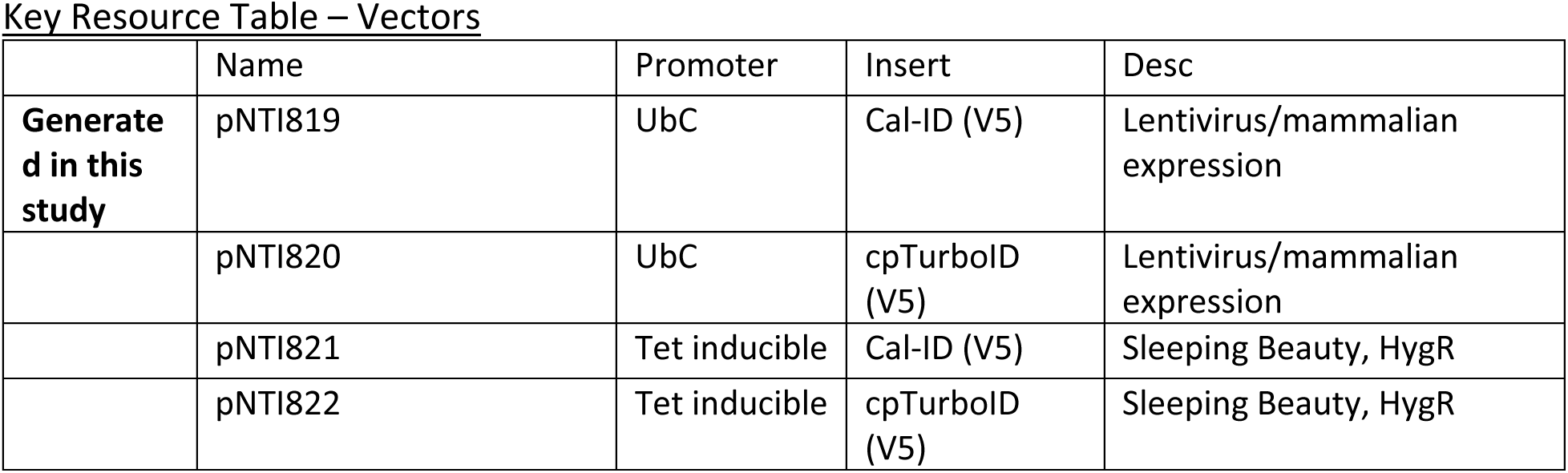
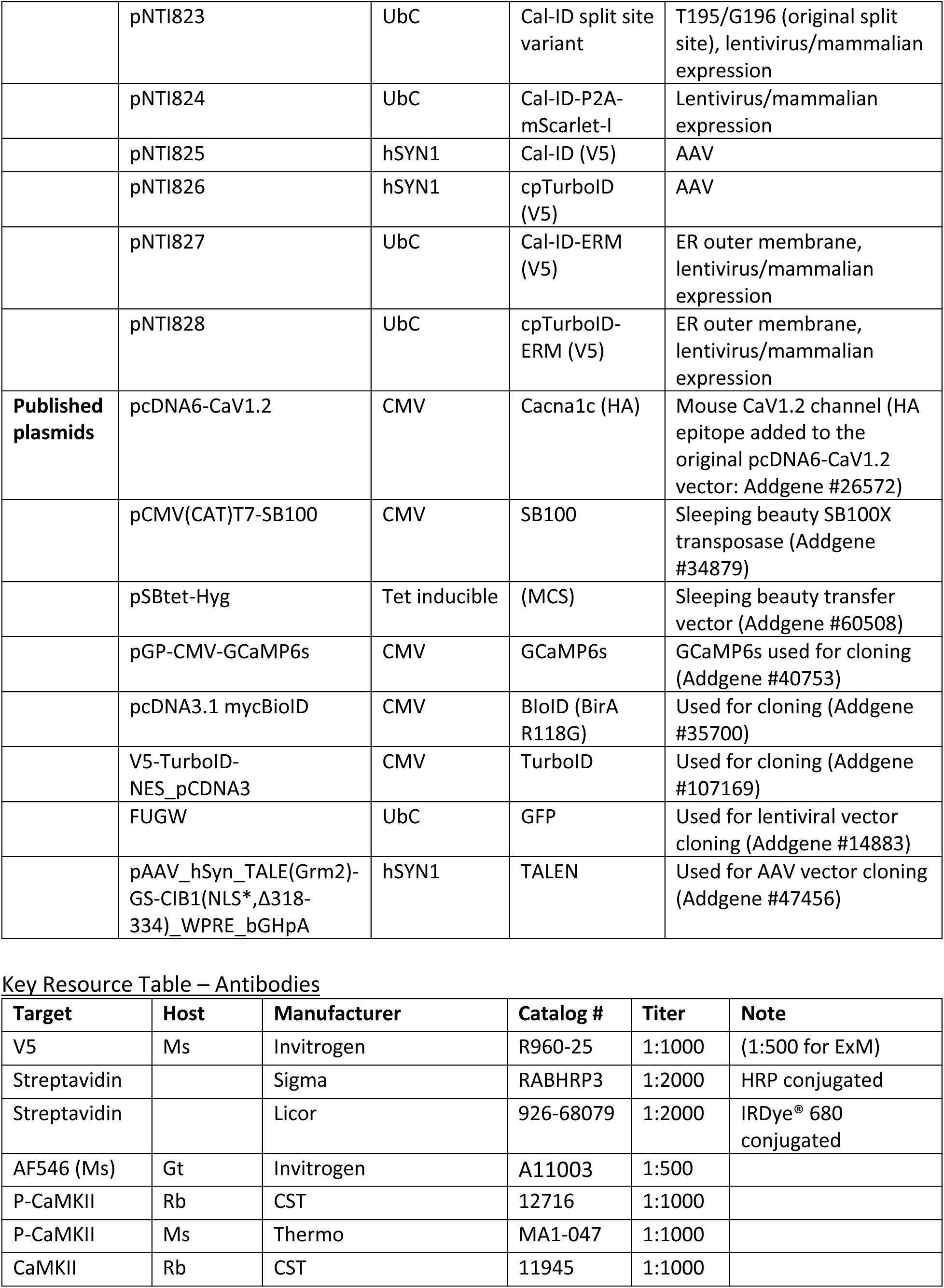
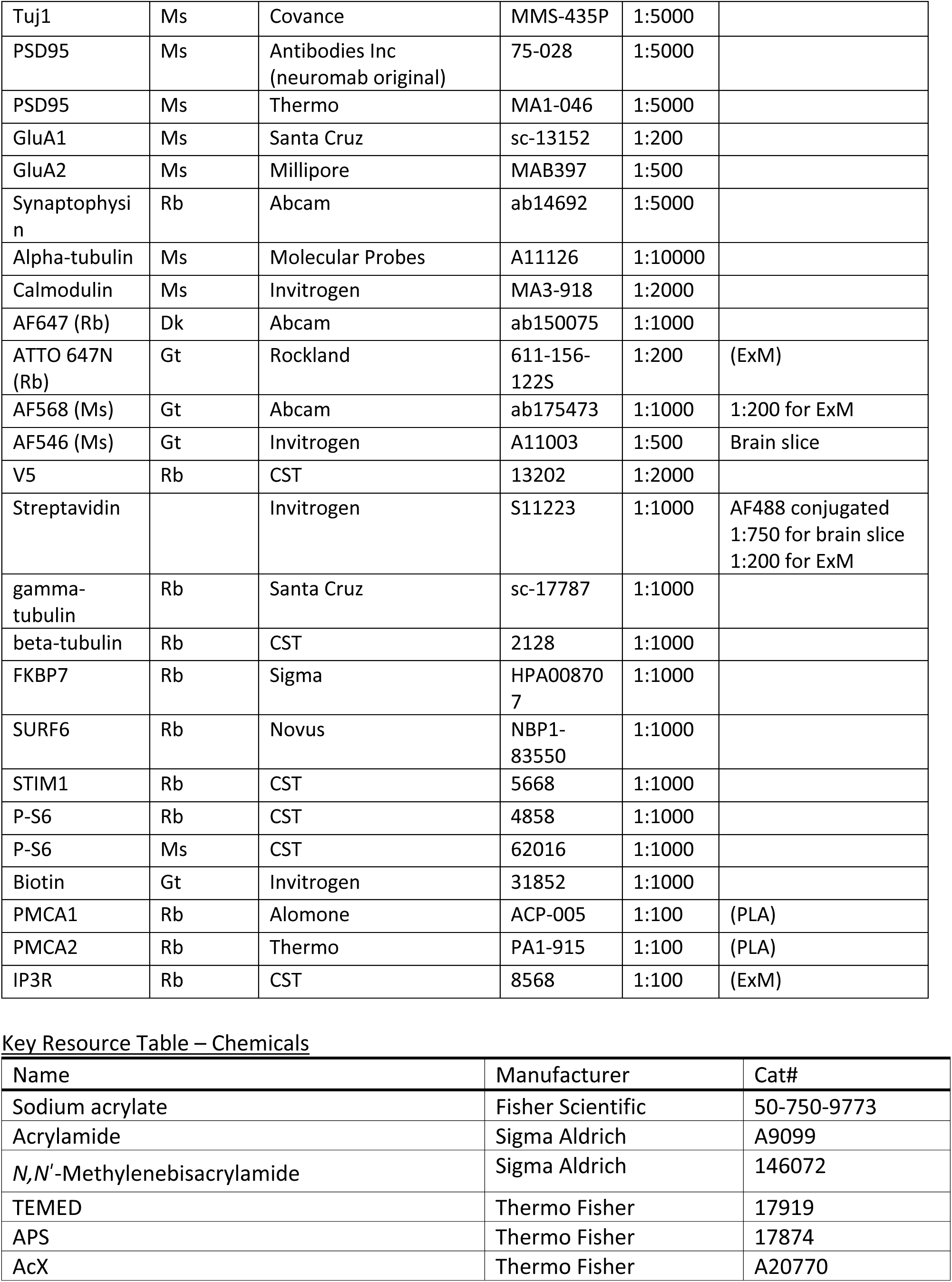
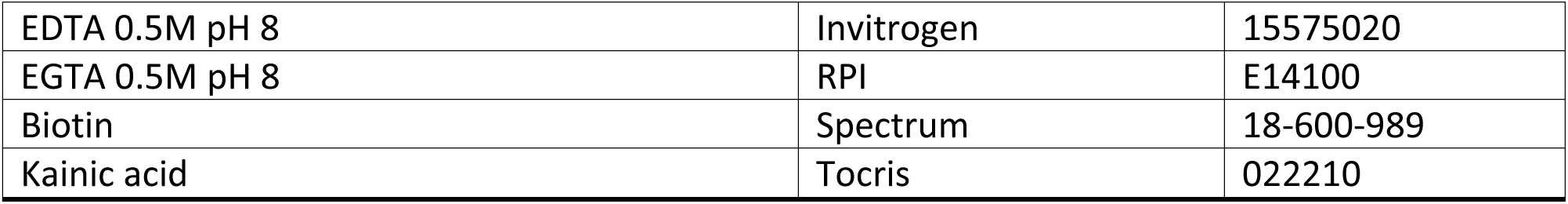
Mass spectrometry and imaging analyses results.

**Supplementary Movies 1 and 2. 3D projection: Cal-ID-ERM/ExM, HEK293T cell.**

A 3D projection movie from a Cal-ID-ERM-labeled and expanded HEK293T cell with V5/IP3R co-staining. Biotin: 100 µM, 20 min incubation post serum replenishment. Movie 1: Green: streptavidin, Red: V5 (Cal-ID-ERM). 17 z-stack images, 3 µm interval. Movie 2: Green: streptavidin, Red: V5 (Cal-ID-ERM), Magenta: IP3R. 9 z-stack images, 3 µm interval.

## Acknowledgements

We thank S. Upadhyayula, S. Eacker, and members of the Ingolia lab for discussion, J. Lobel for biochemistry, M. Farhan for mouse work. Mass spectrometry was conducted at the UC Davis Proteomics Core. Confocal imaging was conducted at the CRL Molecular Imaging Center, RRID:SCR_017852, supported by Gordon and Betty Moore Foundation.

## Author contributions

J.W.K. and N.T.I. conceived the study and designed the experiments. J.W.K., A.J.H.Y., E.E.A. conducted the experiments. J.W.K. and N.T.I. analyzed proteomics data. J.W.K. performed expansion microscopy experiments with help from W.W. and R.G.. Y.N.J. supervised biochemistry and microscopy experiments with neurons. H.S.B. oversaw *in vivo* experiments.

T.M.D. and V.L.D. oversaw conceptualization of the study. J.W.K. and N.T.I. wrote the manuscript with input from all authors.

## Data and materials availability

All data generated or analyzed during this study are included in the manuscript and source data files. Plasmids and libraries will be available through Addgene. Raw mass spectrometry data are available via ProteomeXchange with identifier PXD033244. All source code used for data analysis will be available upon request.

---

**Extended Data Fig. 1.**
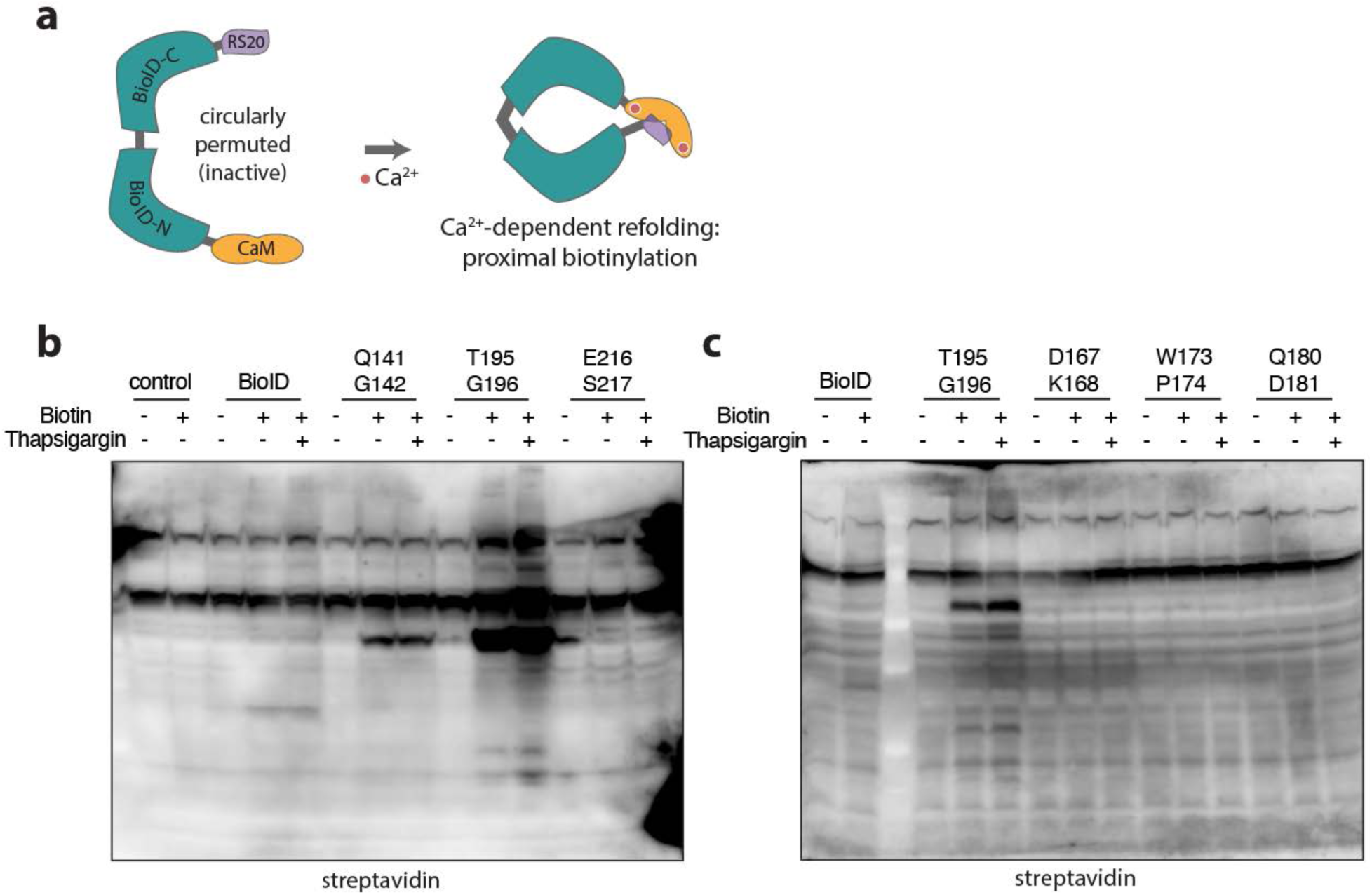
BioID split site screening. (**a**) A schematic diagram of the Cal-ID design used to screen BioID split sites (this early version used split BioID rather than TurboID). CaM: calmodulin, RS20: CaM-binding peptide. (**b)** and (**c**) Cal-ID activation results in HEK293T cells with 1 µM thapsigargin. Biotin: 50 µM, 30 min incubation. Cal-ID constructs were transfected transiently.

**Extended Data Fig. 2.**
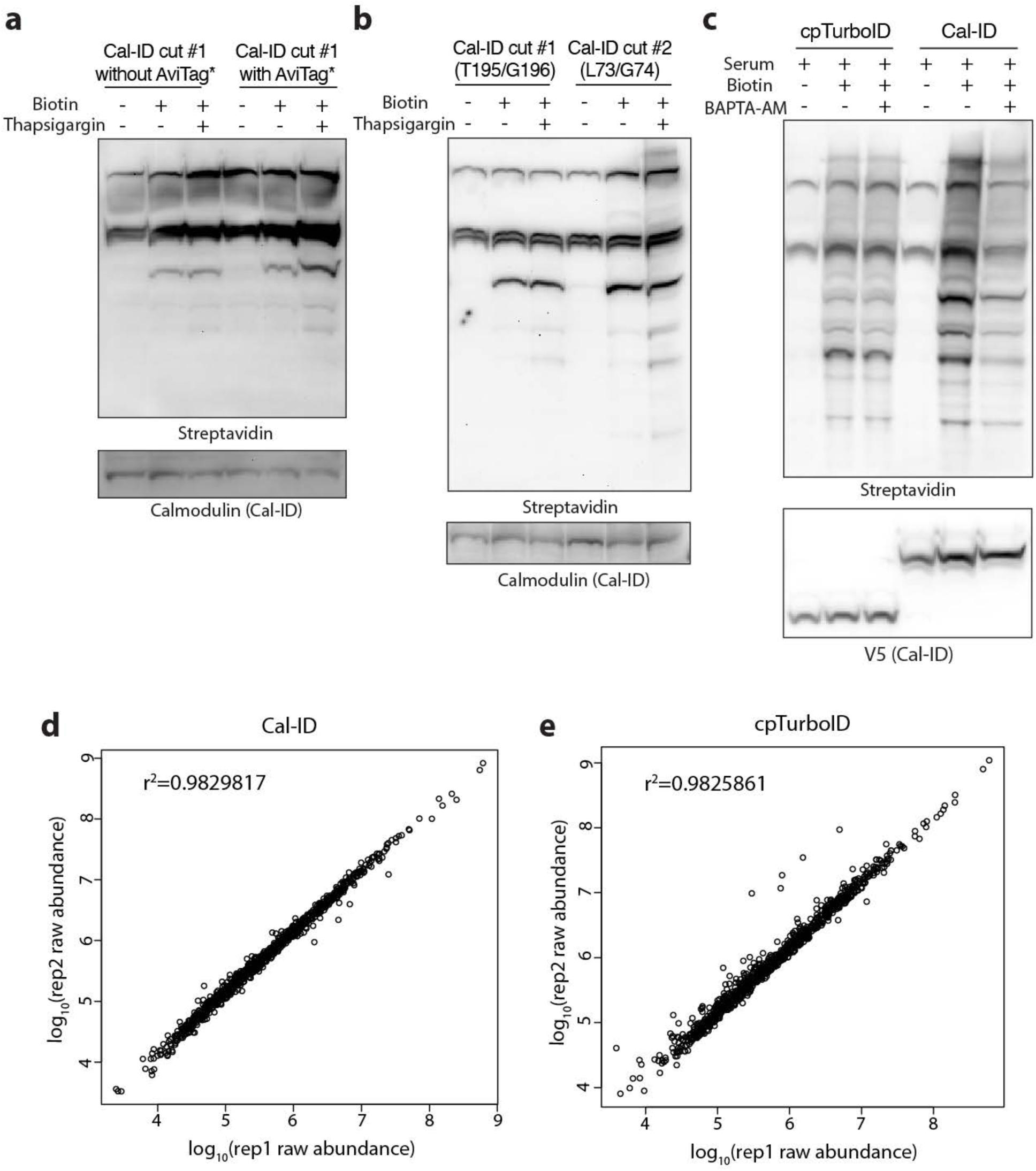
Optimization of Cal-ID and mass spectrometry in HEK293T cells. (**a**) Representative blots testing Cal-ID (T195/G196 split) with a self-inhibitory AviTag* motif in HEK293T cells with 3 µM thapsigargin and prolonged incubation. Biotin: 100 µM, 1 hour incubation, transient transfection. (**b**) Representative blots comparing Cal-ID split site variants T195/G196 and L73/G74 for short incubation. Both variants contain a self-inhibitory motif. Transient transfection in HEK293T cells, 3 µM thapsigargin, biotin: 50 µM, 15 min incubation. (**c**) Representative blots showing that Ca^2+^ chelator BAPTA-AM treatment blocks Cal-ID biotinylation. Transient transfection in HEK293T cells, BAPTA-AM: 40 µM, biotin: 100 µM, serum: replacement with fresh culture media. Culture media was replaced, BAPTA-AM was treated and incubated for 3 min, then biotin was added and labeled for 30 min. V5: cpTurboID or Cal-ID. (**d**) and (**e**) Scatterplots showing mass spectrometry reproducibility between biological replicates (d) Cal-ID, (e) cpTurboID, prepared from HEK293T stable cell lines.

**Extended Data Fig. 3.**
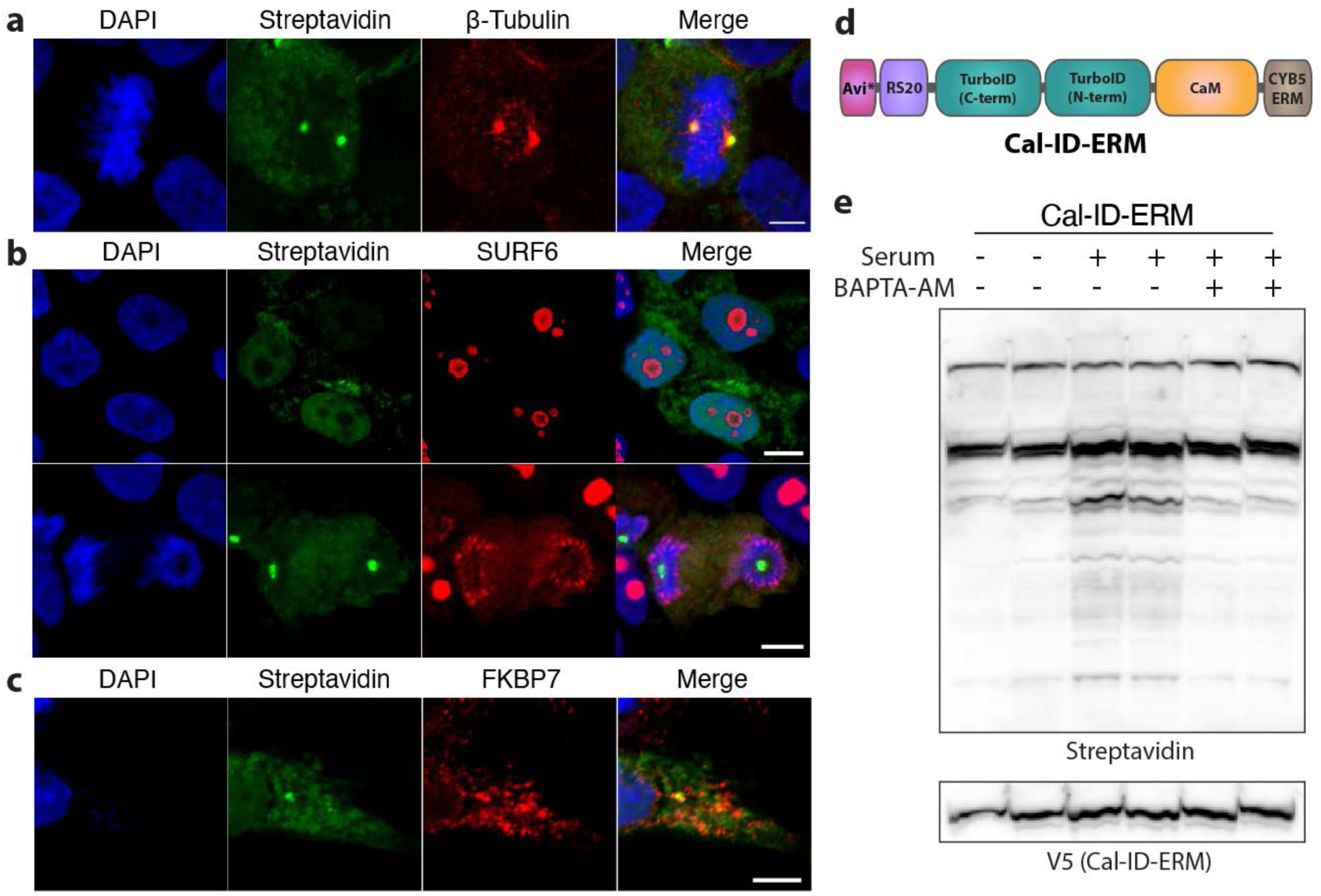
Visualization of Cal-ID target proteins identified by mass spectrometry and ER-localized Cal-ID. (**a** to **c**) Confocal microscope images visualizing biotinylated proteins and mass spectrometry-identified target proteins in Cal-ID expressing HEK293T cells. Co-localization of streptavidin signals and (a) β-tubulin found in mitotic cells, (b) SURF6 in mitotic cells, (c) FKBP7 at the ER. (b) SURF6 did not show co-localization with streptavidin signals during interphase (top) but showed co-localization during mitosis (bottom). Tet-inducible stable HEK293T cells, scale bars = 5 µm. (**d**) A schematic of Cal-ID-ERM, an ER outer membrane-localized Cal-ID. CYB5 ER transmembrane motif is 45 aa long. (**e**) Representative blots showing that Ca^2+^ chelator BAPTA-AM treatment blocks Cal-ID-ERM biotinylation. Transient transfection in HEK293T cells, BAPTA-AM: 40 µM, biotin: 100 µM, serum: 50% culture media replacement with fresh culture media. Ca^2+^ in the DMEM media was chelated using 1.8 mM EGTA in prior to BAPTA-AM treatment. BAPTA-AM was treated and incubated for 3 min, then biotin was added and labeled for 20 min.

**Extended Data Fig. 4.**
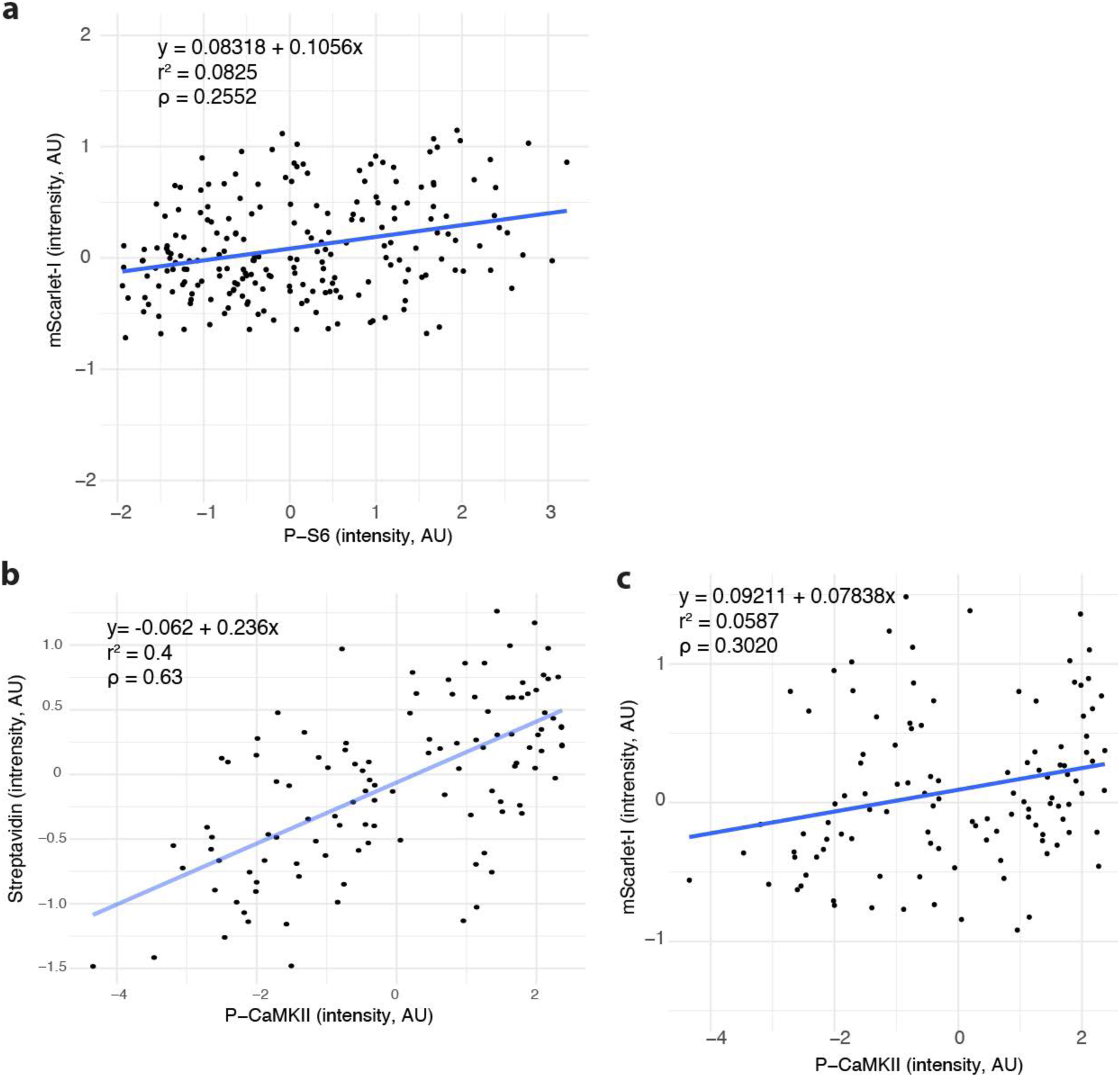
Correlation between neuronal activity markers and Cal-ID biotinylation. (**a**) Scatterplots of P-S6 and mScarlet-I (Cal-ID expression control) signal intensity show a much weaker trend than streptavidin, related to Fig. 4e. (**b** and **c**) Similar to the results from P-S6, P-CaMKII signals show a much stronger positive correlation with streptavidin signal than with mScarlet-I signals. AU: arbitrary unit.

**Extended Data Fig. 5.**
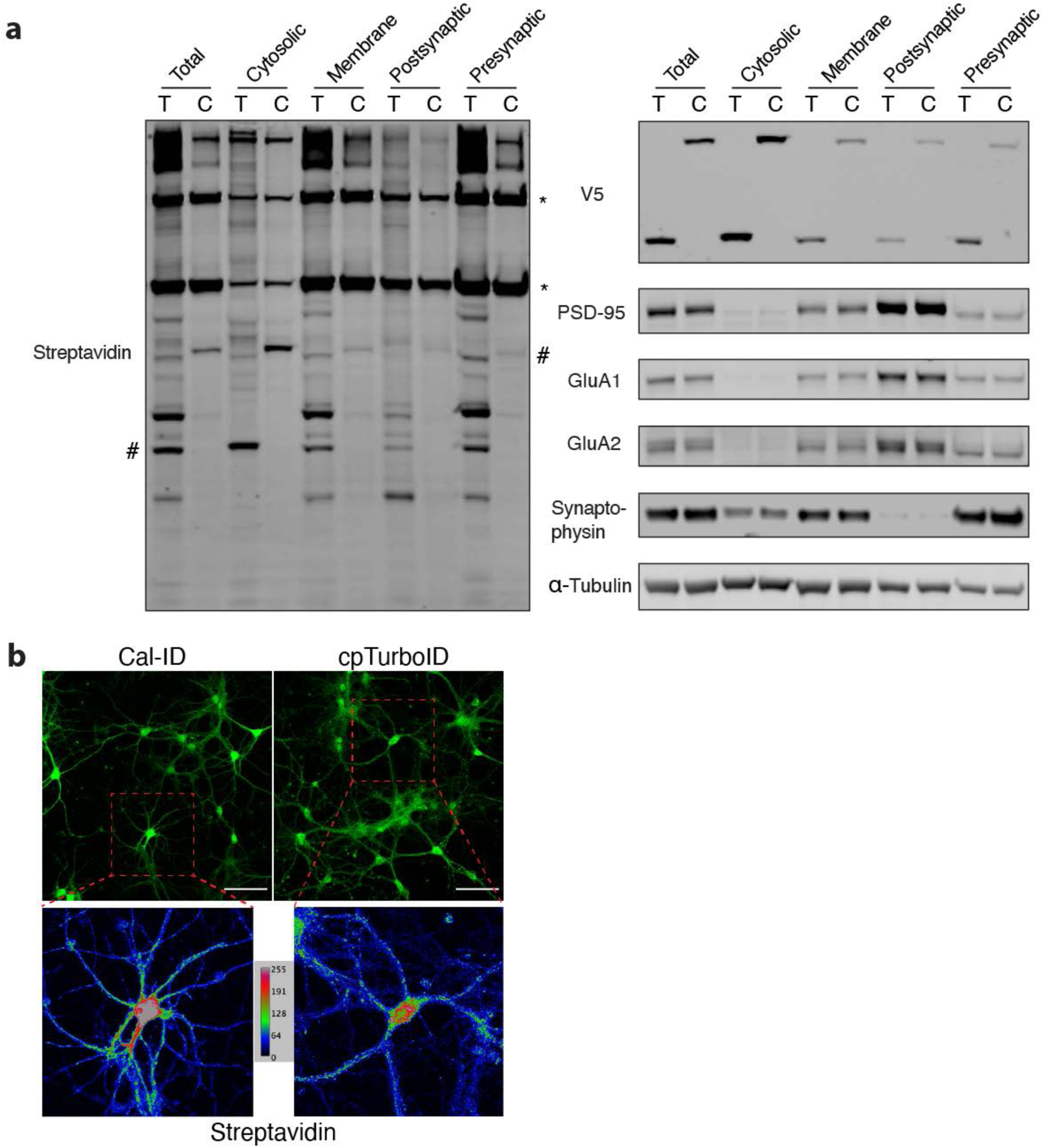
Subcellular localization of Cal-ID in neurons. (**a**) Subcellular fractionation of Cal-ID-biotinylated primary cortical neurons. T: cpTurboID, C: Cal-ID. *: endogenous biotin carrier proteins, #: self-biotinylation of Cal-ID or cpTurboID. PSD-95, GluA1/2: postsynaptic markers, Synaptophysin: presynaptic marker, ⍺-tubulin: loading control, V5: Cal-ID/cpTurboID epitope. biotin: 100 µM, 1 hour incubation. (**b**) Representative images of biotinylation patterns in neurons by Cal-ID or cpTurboID. Color map: signal intensity. Mouse primary hippocampal neurons. Scale bars: 100 µm.

**Extended Data Fig. 6.**
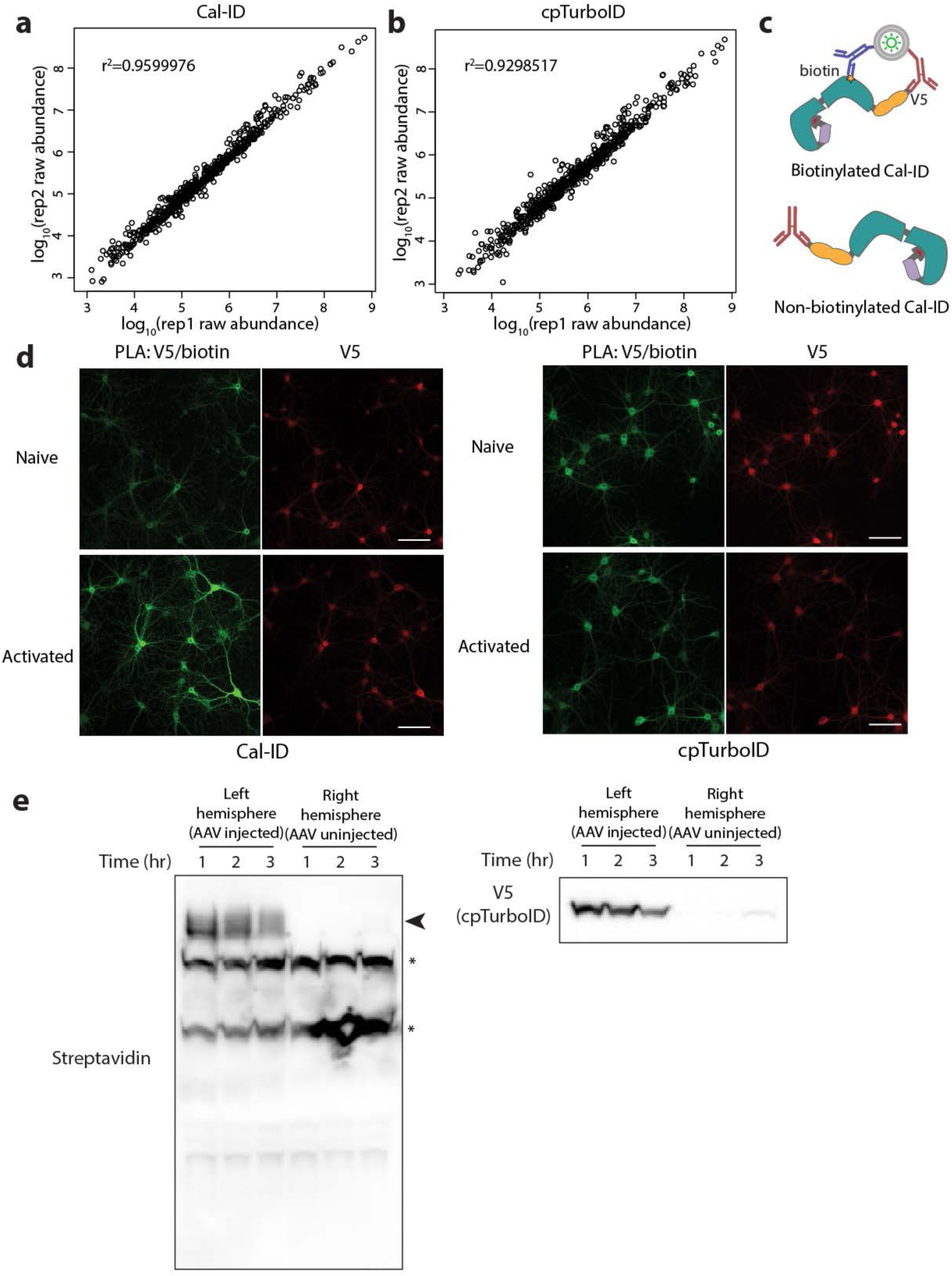
Cal-ID TMT mass spectrometry and proximity ligation assay from mouse primary neurons. (**a** and **b**), Scatterplots showing mass spectrometry reproducibility between biological replicates (a) Cal-ID, (b) cpTurboID, prepared from mouse primary cortical neurons. (**c**) A schematic diagram of proximity ligation assay (PLA) to visualize self-biotinylation of Cal-ID. (**d**) PLA was performed with V5 (Cal-ID) and biotin. Activated: 50 µM APV + 10 µM NBQX for overnight silencing, followed by 4 mM CaCl2 along with 100 µM biotin for 1 hour. Mouse primary hippocampal neurons, lentiviral delivery, scale bar: 100 µm. (**e**) *In vivo* biotinylation time course results. Unilateral (left), intracortical injection of cpTurboID-encoding AAV was performed at P0/1. Biotin injection was performed at 4 weeks of age. Biotin: 24 mg/kg in PBS (i.p.). 1 to 3 hours of post injection incubation time before euthanasia. N = 1 per time point. *: endogenously biotinylated proteins. Arrow: cpTurboID-mediated biotinylation.

**Extended Data Fig. 7.**
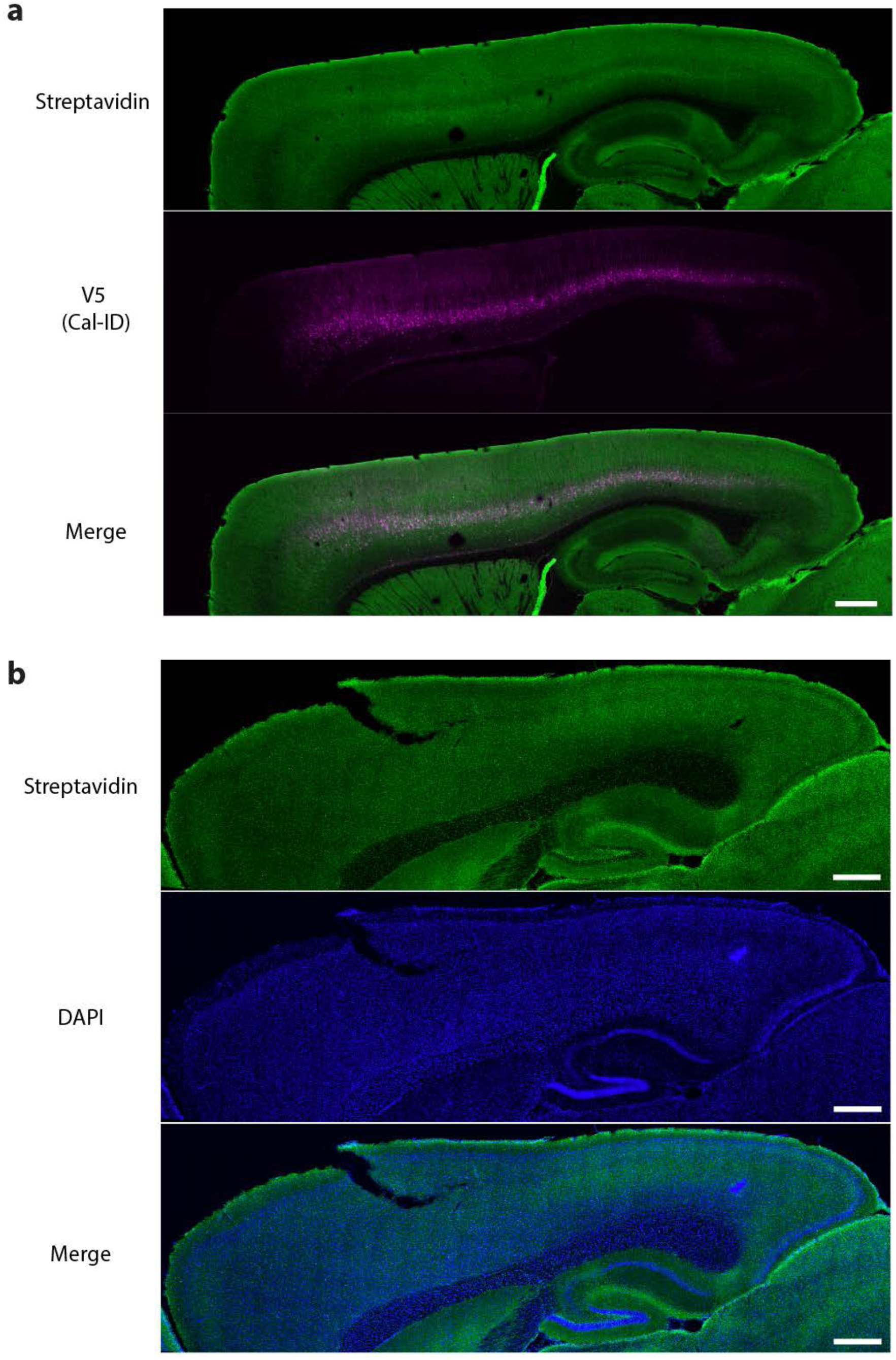
Cal-ID imaging of saline-injected mouse brain. (**a**) Representative images from a brain slice of saline-injected, Cal-ID labeled mouse brain. Sagittal section with 40 µm thickness. Scale bar: 500 µm. Biotin: 24 mg/kg, kainic acid: 15 mg/kg. Biotin intraperitoneal injection was followed by 45 min incubation, then saline was intraperitoneally injected and another 45 min incubation before perfusion. (**b**) A representative image from the mouse cortex without AAV injection; this image is from the right hemisphere (uninjected side) from a mouse received Cal-ID AAV injection to the left hemisphere and was treated with biotin and kainic acid i.p. injections. Scale bar: 500 µm. Biotin: 24 mg/kg, kainic acid: 15 mg/kg. 40 µm thickness sagittal section.

**Extended Data Fig. 8.**
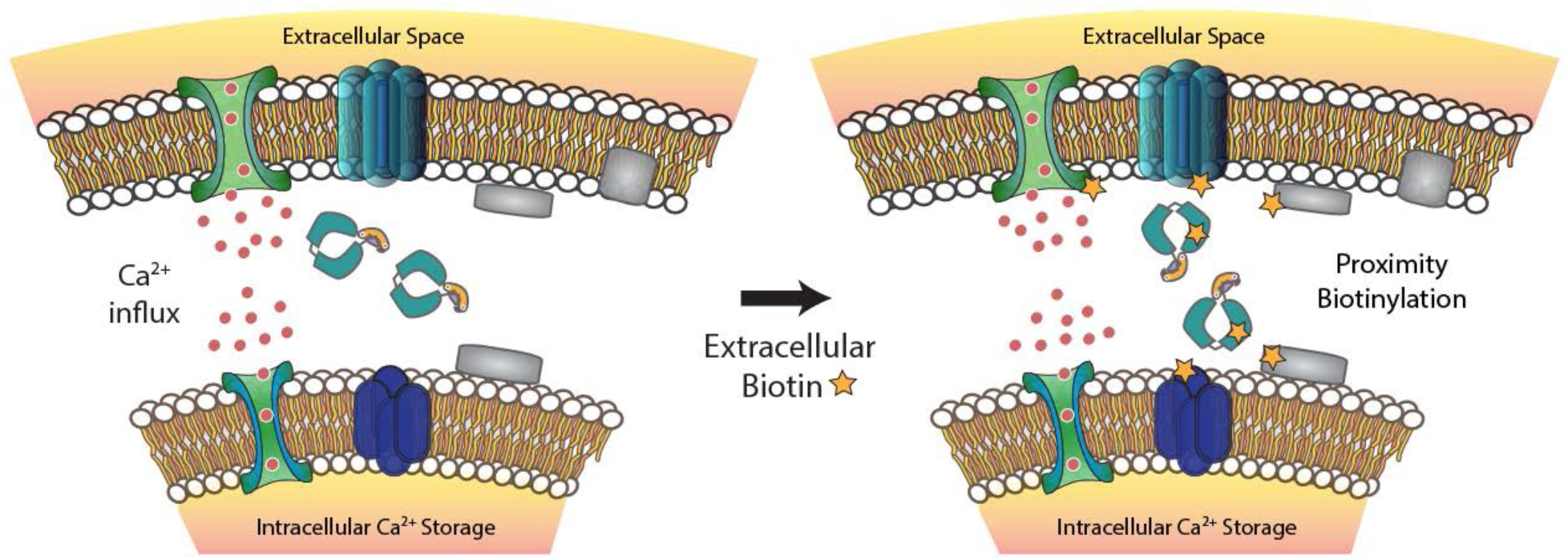
Cal-ID biotinylates calcium signaling microdomains in living cells. A schematic illustration of the action of Cal-ID for the labeling of Ca^2+^ concentration microdomains in living cells.

